# The response time paradox in functional magnetic resonance imaging analyses

**DOI:** 10.1101/2023.02.15.528677

**Authors:** Jeanette Alane Mumford, Patrick G. Bissett, Henry M. Jones, Sunjae Shim, Jaime Ali H. Rios, Russell A. Poldrack

## Abstract

The functional MRI (fMRI) signal is a proxy for an unobservable neuronal signal, and differences in fMRI signals on cognitive tasks are generally interpreted as reflecting differences in the intensity of local neuronal activity. However, changes in either intensity or duration of neuronal activity can yield identical differences in fMRI signals. When conditions differ in response times (RTs), it is thus impossible to determine whether condition differences in fMRI signals are due to differences in the intensity of neuronal activity or to potentially spurious differences in the duration of neuronal activity. The most common fMRI analysis approach ignores RTs, making it difficult to interpret condition differences that could be driven by RTs and/or intensity. Because differences in response time are one of the most important signals of interest for cognitive psychology, nearly every task of interest for fMRI exhibits RT differences across conditions of interest. This results in a paradox, wherein the signal of interest for the psychologist is a potential confound for the fMRI researcher. We review this longstanding problem, and demonstrate that the failure to address RTs in the fMRI time series model can also lead to spurious correlations at the group level related to RTs or other variables of interest, potentially impacting the interpretation of brain-behavior correlations. We propose a simple approach that remedies this problem by including RT in the fMRI time series model. This model separates condition differences from RT differences, retaining power for detection of unconfounded condition differences while also allowing the identification of RT-related activation. We conclude by highlighting the need for further theoretical development regarding the interpretation of fMRI signals and their relationship to response times.

## Introduction

The goal of task-based functional magnetic resonance imaging (fMRI) studies is to infer the involvement of particular brain regions or networks in specific cognitive functions. These studies are most often designed using the subtraction logic first developed by ***Donders*** (***1969***) for the analysis of response times (RTs), in which comparisons are made between different task conditions that are thought to differ with regard to the involvement of some specific cognitive function(s). For example, in the well known Stroop task, stimuli are presented in which the color and text of the word are either congruent (e.g. “blue” presented in blue) or incongruent (e.g. “blue” presented in red) (***Stroop, 1935***). Individuals are consistently slower at naming the color of the stimulus when the written word is incongruent compared to congruent, and this difference in response times is interpreted as indexing the engagement of an additional cognitive process in the incongruent condition, such as conflict detection or resolution (***Botvinick et al., 2001***). Similarly, greater activation in regions such as the dorsal medial frontal cortex (dMFC) in fMRI studies of the Stroop task have been interpreted as reflecting a specific role in these cognitive processes (***Botvinick et al., 1999***; ***MacDonald et al., 2000***; ***Kerns et al., 2004***).

The facile nature of the common inference from activation to “involvement” belies the deep complexity of the link between fMRI signals and underlying neuronal activity (cf. ***Logothetis*** (***2008***)). Here we focus on disambiguating these interpretations of activation: namely, whether a difference in activation reflects the differential engagement of a particular computation, or engagement of the same computation for a different amount of time. Because of the slow nature of the blood oxygen-level dependent (BOLD) response that is measured in most fMRI studies, it is nearly impossible to distinguish the degree to which a difference in evoked BOLD response reflects an increase in the amplitude of neuronal response versus a difference in the duration of that response (Figure 1). This indeterminacy has been known since the early days of fMRI (***Savoy et al., 1995***; ***Jezzard et al., 2001***), and establishes the importance of considering the potential of differences in the duration of neural activity to confound amplitude estimates in some fMRI tasks.

**Figure 1.**
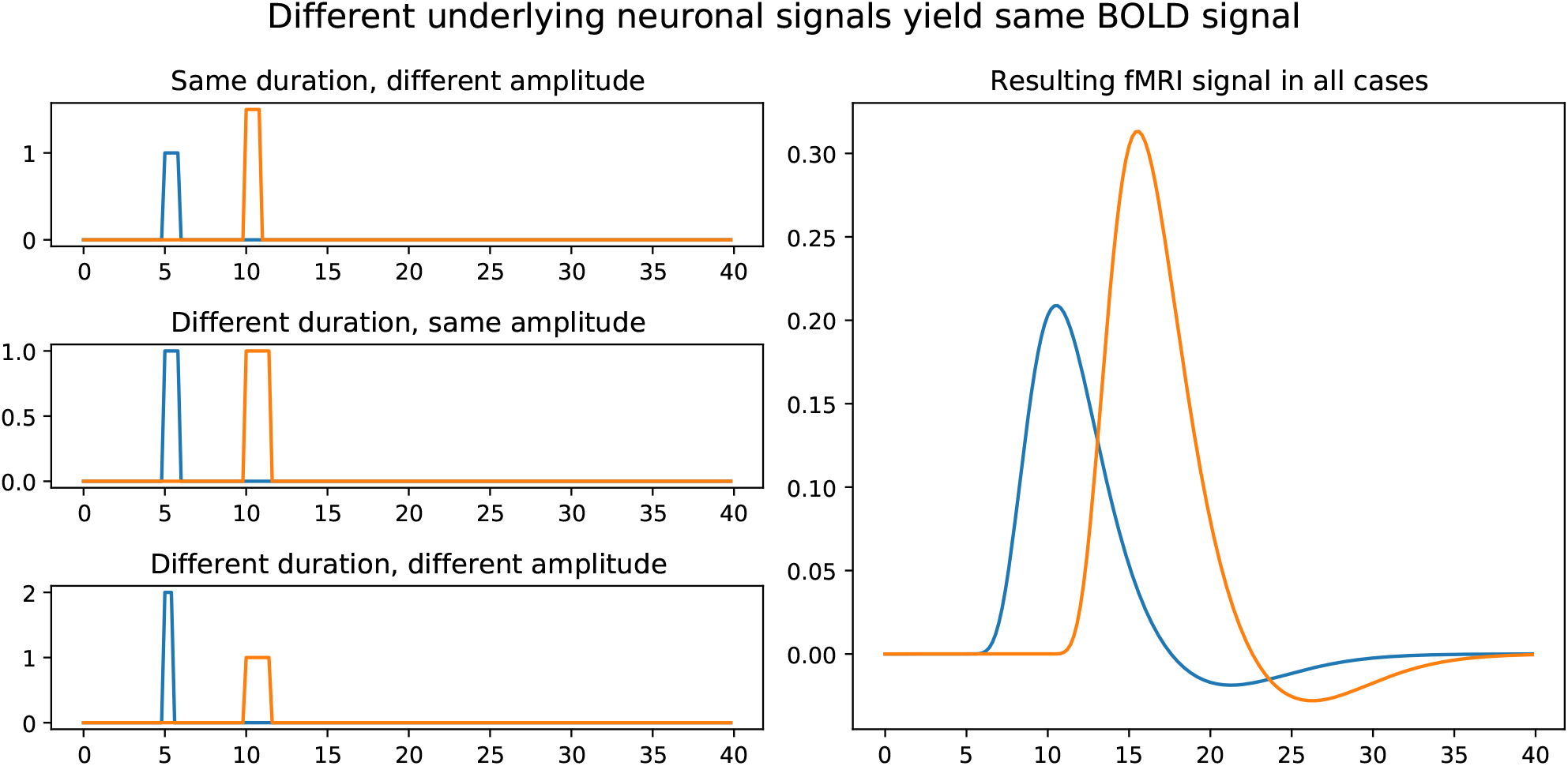
Illustration of how amplitude and duration of neuronal signal interact to yield similar BOLD responses. The left hand column shows 3 different examples of neuronal signals that evoke the same BOLD response shown in the right hand panel.

Over a decade ago, ***Grinband et al.*** (***2008***) started the discussion about modeling of response times in fMRI analysis. They proposed the use of a variable epoch model, where the trial-by-trial neuronal durations were assumed to track with RTs as opposed to the common modeling practice at the time that assumed each trial was sufficiently modeled as a brief impulse of constant duration (e.g., .1s). The impulse model was previously considered to be adequate in rapid event-related designs due to the belief that differences in RTs would not be detectable in this setting. The proposed model using RTs as the duration was shown to produce activation estimates that were not confounded by RT differences and yielded more powerful results than the impulse model or an impulse model that included an RT-modulated regressor. Importantly, this model’s performance studied within-subject power for a single condition versus baseline and relied on specific assumptions about the underlying signal, specifically that the duration of neuronal activation mirrors the RTs, which is not necessarily the case. The behavior of this model when this assumption is not met has not been well studied nor has the behavior of more commonly used condition difference contrasts in the context of response time differences between conditions.

***Yarkoni et al.*** (***2009***) examined the relationship between RT and fMRI signal across a variety of tasks in an effort to better understand how RTs were related to the measured BOLD signal. Evidence was found that RT-driven amplitude differences were likely due to “time on task” or simply duration differences in the neuronal signal across trials, as opposed to differences in the magnitude of neuronal activity (characterized as “effort” in that paper). These time on task effects could reflect stimulus-related processes that simply reflect constant neuronal activity that varies in relation to the duration of the trial, rather than differences in the amplitude of neuronal activity across trials. A compelling result of this work was the identification of a widespread network of brain regions that showed significant correlation with RTs across each of a variety of different tasks including the dMFC which was previously described as reflecting conflict in the Stroop task. This calls into question how RT-based activation differences might be interpreted given that they are common across many tasks and thus are unlikely to reflect processes that are specific to a particular task.

A subsequent series of papers focused on the incongruent versus congruent contrast of the Stroop task and whether activation in the dMFC reflected conflict, as proposed by a prominent theory (***Botvinick et al., 2001***). This inspiring discourse across multiple publications illustrates the challenge of interpreting RT-correlated activation as well as demonstrating the rigorous work required to combine the behavioral theory of a task with imaging analysis results when RT-correlated activation is found. Both ***Grinband et al.*** (***2011b***) and ***Carp et al.*** (***2010***) showed that the difference in activation between slow and fast congruent trials was similar to the difference between all congruent and incongruent trials in the dMFC, supporting the idea that the commonly found effect was driven by RTs. The next step to fully understand this RT-driven effect is the difficult step of determining whether this is a time on task effect or an actual difference in the amplitude of neuronal signaling that correlates with RT and whether these differences align with the underlying theory of conflict in the Stroop task. This was the focus of follow-up work by ***Yeung et al.*** (***2011***) who proposed that the RT-based effects were a result of differential engagement of specific cognitive processes and not simply due to time on task. They also argued that the result supported the predictions from the computational model that formalizes the conflict monitoring theory. Even though this series of papers drew attention to how we model and interpret response times in the fMRI based Stroop task, a consensus was not reached (***Brown, 2011***; ***Grinband et al., 2011a***; ***Nachev, 2011***) and it did not have a widespread impact on how the Stroop task is modeled or interpreted in fMRI data. Of the 22 papers published resulting from a PubMed search for “stroop task fmri 2021”, only 4 addressed RT in their analyses and interpretation of their results.

It is beyond the scope of the present work to come to an agreement on how to interpret Stroop-based fMRI activation maps that may be driven by RTs, but we use this example to illustrate challenges of modeling RT-based effects in fMRI results and the important implications it can have on theoretical conclusions. If a model only evaluates condition differences, without adjusting for RTs, an observed condition effect has multiple potential interpretations, as the true effect could be: a simple condition difference in the amplitude of neuronal response (not driven by RTs), an RT effect with constant amplitude of the neuronal response (with no condition difference) or a relationship where there is both a condition difference and RT effect. If the result is driven by RTs the question remains as to whether it reflects time on task versus a difference in the amplitude of neuronal signaling. Therefore, our ability to interpret the finding of an unadjusted condition difference, without further analysis and theoretical work, is limited at best.

For clarity, we define some terms that will be used throughout this paper. Time series-level analysis refers to linear models of fMRI time series. A group-level analysis refers to an analysis of an estimated fMRI contrast across a set of subjects that could be a single group average (1-sample t-test), group average comparisons (e.g., 2-sample t-test), linear associations with a covariate (e.g., phenotype) or other group-level models. Between-trial RT adjustment is formally carried out in the time series-level. Between-subject RT confounds will only impact group-level models involving group comparisons or associations but not single group averages. Although we focus on “2 stage” models, with only within-subject and group levels, these ideas extend to three stage models where subjects have multiple runs and so analyses are done within-run (time series analysis), within-subject (combining runs) and between-subject.

Limited focus has been given to the link connecting between-trial RT adjustment in the time series analysis to between-subject RT confounds in the group-level analysis. For example, if between-trial RT adjustment is ignored, the incongruent versus congruent Stroop contrast may be correlated with the within-subject difference in average RT between incongruent and congruent trials. In earlier work (***Carp et al., 2012***), this relationship was illustrated by showing that the correlation between age and the contrast estimate of incongruent versus congruent Stroop conditions changes based on whether between-trial RT adjustment is performed. Although this work points out the possibility of between-trial RT adjustment impacting between-subject analyses, the link has not been formally defined.

In the present work we extend and improve upon previous attempts to incorporate RT into in fMRI data analysis. We present a model that separates condition differences, adjusted for RTs, and RT-specific effects. This model is flexible, obviating the assumption in ***Grinband et al.*** (***2008***) that the duration of the neuronal signal necessarily matches the RTs. When the duration of the signal matches RTs, we refer to this type of BOLD signal as “scaling with RT”. The RT duration model can bias results when the underlying true signal duration is constant across trials (i.e., does not scale with RT), whereas the model presented here can adapt to either of these situations. We further quantify the bias in the ***Grinband et al.*** (***2008***) model as well as the most commonly used model that ignores RTs completely.

We also formally define the relationship between the between-trial RT adjustment, or lack thereof, and potential between-subject RT confounds. We will show that when between-trial RT adjustment is skipped, the effect size magnitude of these potential between-subject RT confounds is on the order of common effects of interest, e.g., correlations with various behavioral phenotypes. An unexpected, but concerning result occurs if a variable (e.g., age) is associated with each condition, separately, but this variable is not associated with the condition difference. If RT is ignored in the time series analysis it can introduce both an RT effect and a false association with the variable (age in this example), and the only way to remedy this problem is to repeat the time series analysis with a model that adjusts for between-trial RT differences. This will be a barrier for some when using condition difference estimates supplied in databases for large neuroimaging studies, since RT adjustment in the times series model is typically skipped and redoing the time series analyses for thousands of subjects may not be possible for many users of these databases.

Finally we replicate and extend the findings of ***Yarkoni et al.*** (***2009***) by demonstrating that the widespread association between RT and fMRI activation is consistent over a set of 7 fMRI tasks, each with approximately 91 subjects.

The overarching goal of this work is to revive an interest and curiosity in understanding and addressing RT-correlated activation to improve our interpretation of fMRI signals and their relationship to underlying theories of RT-based tasks. The suggestions we present are easy to implement and enable the direct estimation of RT-based effects in parallel with condition differences, adjusted for RTs. This does not limit the researcher, but instead more clearly defines what is being studied, thus providing a cleaner link to underlying behavioral theory and improving the ability of fMRI to help understand brain function.

## Results

### Simulations

The statistical models used for fMRI data generally involve the convolution of a vector representing trial or stimulus onsets with a canonical hemodynamic response function (as shown in Figure 1) to create regressors for use in linear modeling (***Poldrack et al., 2009***). The trials can be represented either as delta functions or as boxcar functions with some duration; when a boxcar is used, it is common to set the duration of the boxcar to a constant value such as the duration of the stimulus, or to some brief default value (such as 0.1 second). In Figure 2 this model is referred to as “Constant duration, no RT” (hereafter as ConstDurNoRT). Because of the indeterminacy described above (Figure 1), the specific constant value used for stimulus duration will not generally impact the statistical inferences derived from the model, as it will simply scale the values of the parameter estimates along with their variances (assuming the trial durations are relatively short). This standard approach does not include any information about response times; thus, if two conditions differ in their RTs when the true activation magnitude does not differ, the condition with the longer RTs may have higher estimated activation if the signal scales with RT.

**Figure 2.**
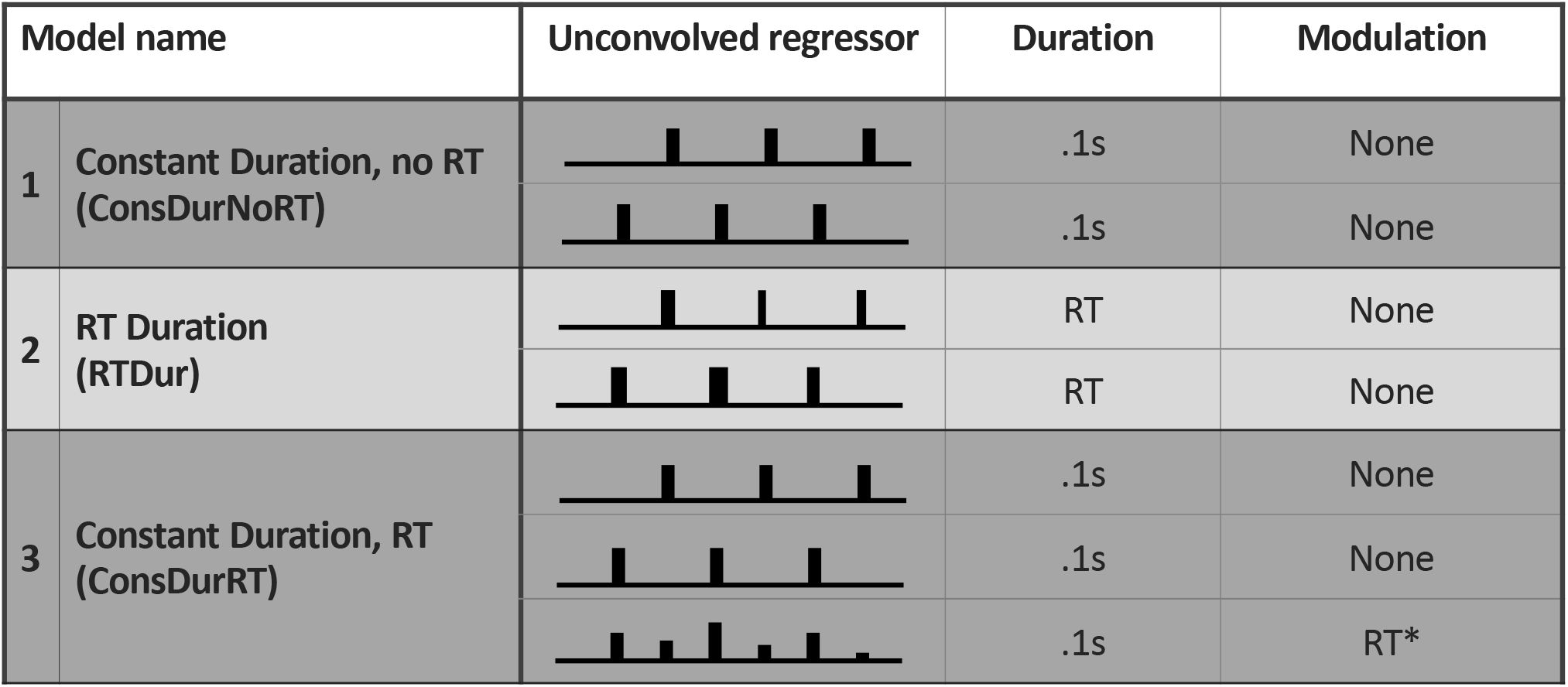
Models assessed in the simulation study described by name, unconvolved regressor visualization, duration used for boxcars of unconvolved regressors and definition of the modulation used, when present. Convolved regressors were used in data generation and modeling. The first model does not include any response time information, the second model addresses RT through the duration of the regressors and the third model adds an RT modulated regressor to the first model. *See Discussion for details on why RT is not centered and other details about centering.

***Grinband et al.*** (***2008***) developed a modeling approach to address the confounding effect of response times in fMRI data, in which the duration of the boxcar function for each trial was varied by the response time on that trial (labeled as “RT Duration” in Figure 2 and hereafter as RTDur). This approach will appropriately scale the parameter estimates for regions in which neural activity duration matches the RT duration, which we will refer to as “the signal scales with RT”. Technically, this model implies a restricted condition by RT interaction model, in that it is modeling a separate RT slope, by condition, without main condition effects (i.e., constant duration regressors for each condition). Specifically, the model implies each condition has a different linear relationship between BOLD activation and RT, but the intercepts (BOLD activation when RT is 0) are both zero. Given this, the model has two shortcomings. First, it will not correctly model activation in regions where neural activity does *not* scale with RTs but rather the true duration is an unknown constant across all trials. Second, it does not allow a separate identification of condition differences versus RT effects; instead, it only performs well if the restricted interaction model is correct.

To address these issues, we created a generalized model of RT that can identify RT effects separately from the task effect (corrected for RT); this is shown as “Constant Duration, RT” in Figure 2 and hereafter as ConstDurRT. This model includes a boxcar function with constant duration for each of the task conditions, along with a single regressor that models the parametric modulation of the response in relation to RT for each trial. Because all RTs are modeled within a single regressor, any differences in RT between conditions will be removed by this regressor, leaving the condition difference effects to be interpreted as unconfounded estimates of activation in relation to the experimental manipulation. This model can be extended to a full interaction model, if an interaction is suspected, and this will be further described in the Discussion section (“Condition by RT interaction models”) as well as concerns in using an interaction model to study condition differences if there is not a significant interaction present. We focus on a simpler non-interaction scenario in the simulation studies to illustrate the results of interest and these results are nontrivial to extend to the interaction model setting.

Notably we have not mean centered RT or subtracted any value from RT on each trial. This will not have any impact on the estimate of the contrast of interest (condition difference) and would only impact the condition versus baseline contrast estimate in this model. If RT is mean centered, within run, the interpretation of some contrasts becomes, “BOLD activation difference when RT is the mean RT for this run”, necessarily introducing an RT-based confound if this contrast is used as the dependent variable in higher level analyses. More details about the impact of centering RTs are included in the Discussion (“Should the RT modulation values be centered?”), including examples where it is necessary to center in some way and how to do so without introducing a new confound. We also discuss why an RT duration regressor is not used instead of the RT modulated regressor. For all models the effect of interest was the subtraction of condition 2 - condition 1, so the RT modulated regressor in ConstDurRT simply serves as a nuisance regressor to pick up RT variability, when present. Of course if there is interest in the RT effect, the parameter of the RT regressor can also be analyzed, although that is not the focus of the simulation analyses and will be the focus of the real data analysis. Further details regarding the modeling approach can be found in Methods; code and data for all analyses are shared at https://github.com/jmumford/rt_simulations (simulations) and https://github.com/jmumford/rt_data_analysis (real data analysis).

Response time data were simulated based on RTs from two different tasks: the Stroop task (based on real data analyzed below) and reported RT distribution parameters from a Forced Choice task used by ***Grinband et al.*** (***2008***). In each case RTs were generated by sampling from an ex-Gaussian distribution (***Ratcliff and Murdock, 1976***); the specified ex-Gaussian parameters led to RTs that were generally longer for the Forced Choice task (mean = 1337, sd = 706.5) compared to the Stroop (mean = 690, sd = 177.5). Another difference is that the variance relative to the mean is smaller for the Stroop task (coefficient of variation of .528 and .257 for the Forced Choice and Stroop tasks, respectively). The interstimulus interval (ISI) was sampled from a Uniform distribution. Trials were either randomly presented conditions or blocked conditions, where 4 trials of the same condition were presented in a row. Time series data that scale with RT were based off of the RTDur regressors and data that did not scale with RT were generated using the ConstDurNoRT regressors.

All simulation-based results correspond to group level analyses with 100 subjects. See the Methods for further details on effect size and variance settings.

### Error rates and power

We first assessed the false positive rate for each of the models on each of the simulated data sets for the condition comparison contrast (Figures 3, S1). In all cases the ConstDurRT model appropriately controlled Type I error, but error rates were inflated when model assumptions regarding the relationship between RT and neural activity were violated by the data. Specifically, ConstDurNoRT had highly inflated error rates when activation did scale with RT, and RTDur had inflated error rates when the signal did not scale with RT. Thus, the most commonly used model for task fMRI analysis, ConstDurNoRT, suffers from substantial inflation of false positives in the face of RT differences between conditions, because it inaccurately attributes the confounding RT signal to differences in the intensity of the underlying neuronal signal. The larger Type I error rates observed with the Stroop-based RT reflects that the standard deviation of the RT, relative to the mean, was lower in this setting and so RT-based differences are easier to detect. The error rates for blocked designs (solid lines) are slightly higher, likely due to the fact that blocked designs have a higher signal to noise ratio, making it easier to detect RT differences in the data. Results with a longer ISI, between 3-6s, are similar (Results shown in Figure S1).

**Figure 3.**
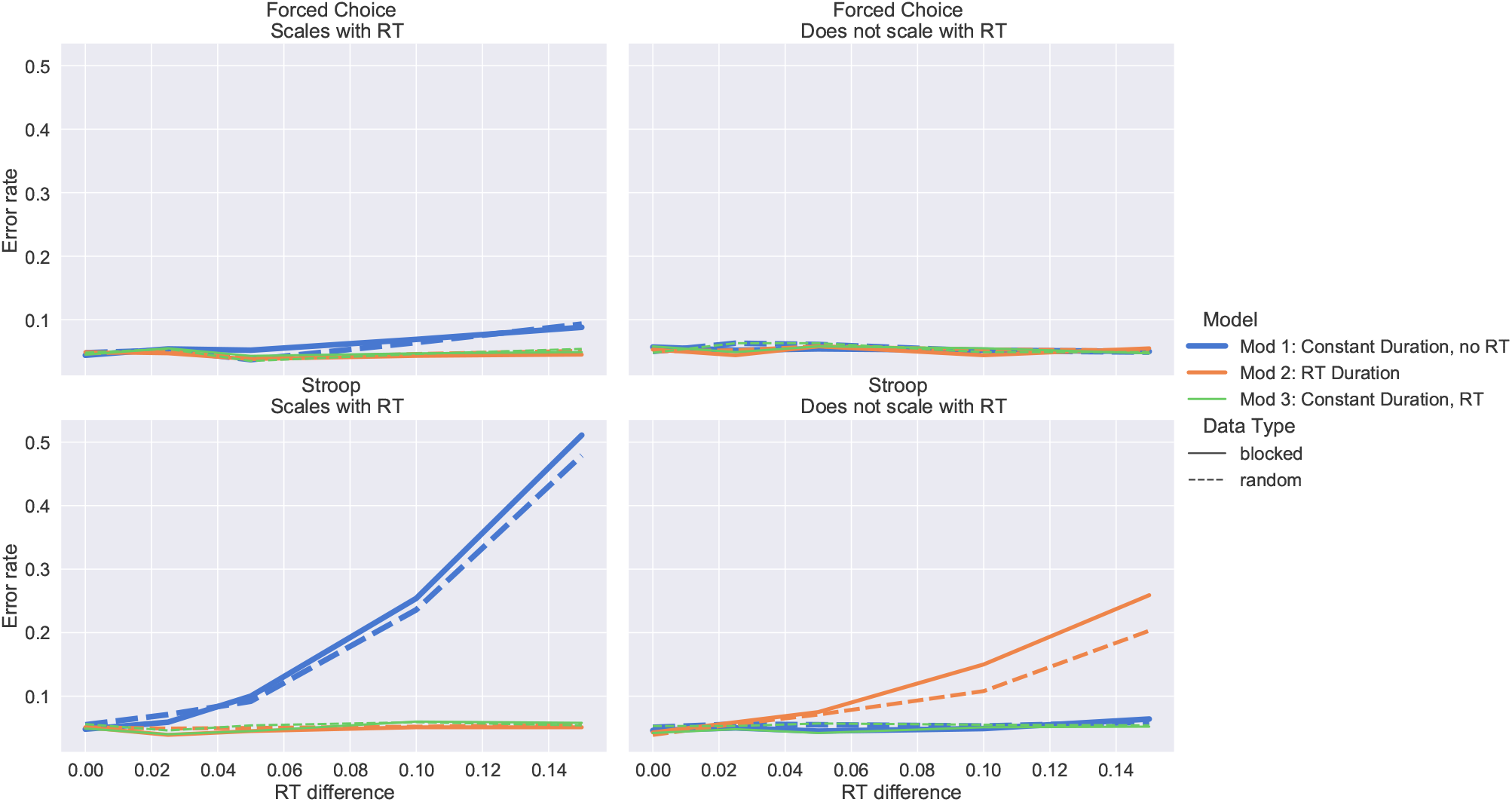
Type I error as RT difference between conditions increases. The Forced Choice Task RT distribution was used in the top panels, while Stroop RT distribution was used in the bottom panels, both with an ISI between 2-4s was used and inference of interest was the 1-sample t-test of the condition effect with 100 subjects. 2500 simulations were used to calculate the error rate.

When signal does not scale with RT, power can only be considered for the ConsDurNoRT and ConsDurRT models since the RTDur model did not have controlled error rates. Figure 4 shows that adding an RT-based regressor, when no RT effect is present, does not impact power at the group level. Thus the flexibility of the ConsDurRT model to adapt to either the scales with or doesn’t scale with RT scenario does not result in a loss in power, which was previously implied in ***Grinband et al.*** (***2008***), which only studied power at the single subject level for a condition versus baseline contrast and not the group level condition difference effect as done here. We do not study power in the context of when the signal scales with RT, since the signal generated by scaling each of the RTDur model regressors by different values would represent an interaction model, where the linear relationship between RT and BOLD differs by condition. In this case the ConsDurRT model is clearly the incorrect model and power is irrelevant. The more suitable model would fit an interaction effect by replacing the single RT modulated regressor in the ConsDurRT model by condition-specific RT modulated regressors. Interaction models will be a topic in the Discussion section (“Condition by RT interaction models”).

**Figure 4.**
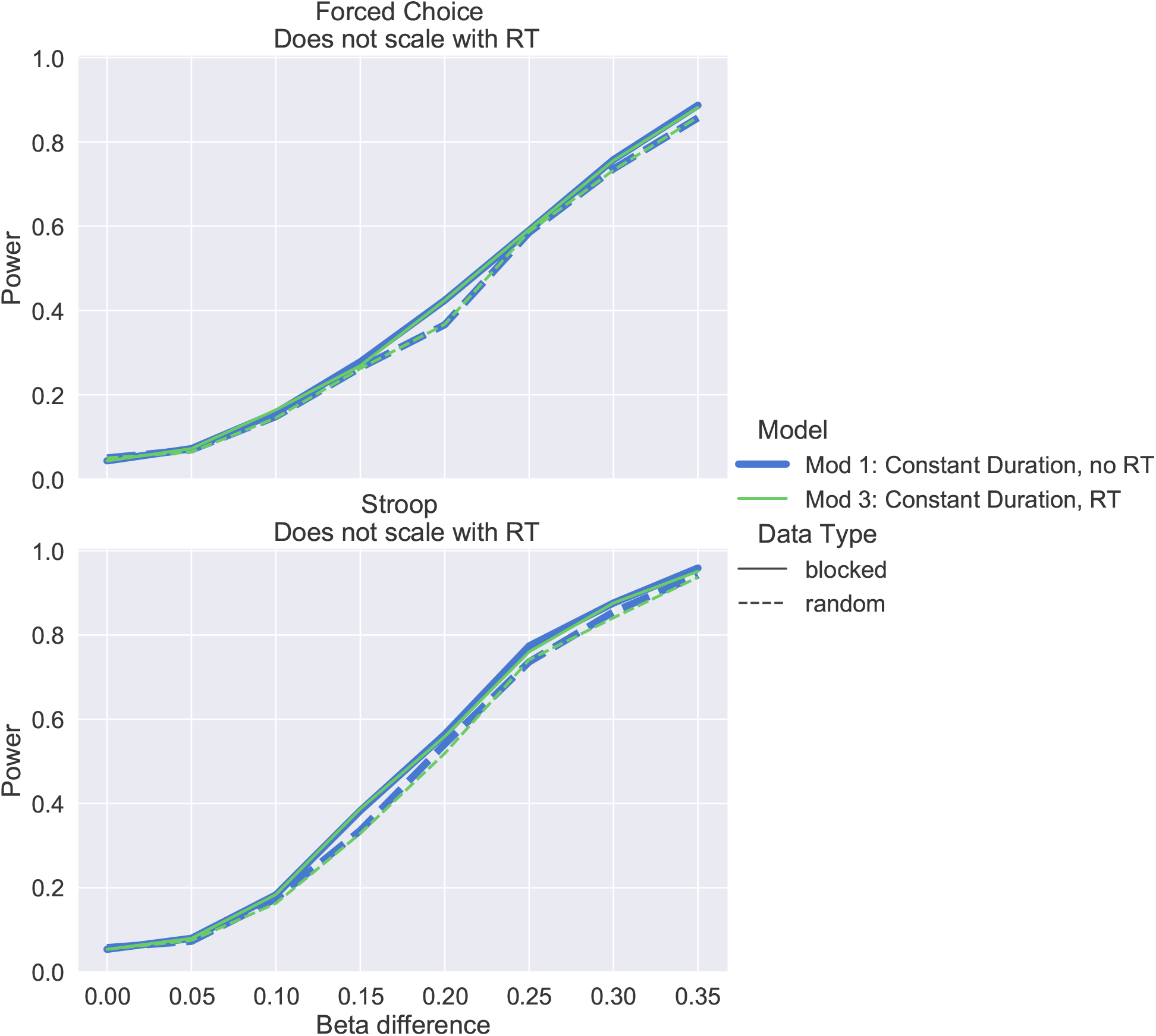
Power when RT difference is 0.1s as the condition difference increases when signal does not scale with RT. The ConsDurRT model (green line) has similar power to the true model, ConsDurNoRT, illustrating no power loss occurs due to including an RT regressor in the time series analysis. Results for RTDur are not shown since it did not have controlled error rates. Sample size is 100, with an ISI between 2-4s.

### RT differences can confound group-level analyses and introduce other false associations

The foregoing analyses, along with the previous work by Grinband, focused on confounding of RT between-trials, which impacts average condition effects. Here we introduce a new problem of a between-*subject* RT confound. The within-subject differences in average RT, corresponding to the contrasted conditions, can confound group level analyses involving group comparisons or associations. For example, the incongruent versus congruent BOLD contrast estimate may correlate with the differences in average RTs for incongruent and congruent conditions. This is of particular interest given the increasing focus on analyses of brain-behavior correlations in fMRI literature (e.g., ***Dubois and Adolphs*** (***2016***)).

The driving factor of correlations between condition differences in brain activation and the corresponding differences in RTs is simply due to a linear relationship between the activation estimate and RT when the data and model assumptions are in conflict. In the case where signals scales with RT and the ConstDurNoRT model is used (duration = 1s), the relationship between the estimated activation, 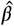, and the true activation, *B*, is approximately 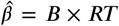, for a single trial. In this illustration, the true activation, *B*, is assumed to be the same for both conditions and does not vary over subjects. Figure 5 shows this linear relationship holds within the range of RTs one would expect to observe in most data sets (i.e., < 2s). Moving from the activation estimate of a single trial to the BOLD activation across multiple trials, the relationship becomes 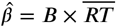, where 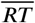 is the average RT across trials. Last, for two conditions, 1 and 2, the relationship for the contrast of conditions is

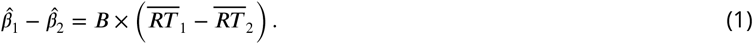

From this it directly follows that in a *group* level analysis there is an expected linear relationship between the estimated condition difference and the difference in RTs, specifically the between-subject slope would be *B*, the true, common activation of the two conditions that does not vary across subjects. As is the case with all linear trends (equivalently, correlations) this relationship does not require a non-zero RT difference on average, but is driven by between-subject RT variability. Therefore the RT difference is an important confound regardless of whether it is significantly different from 0 on average.

**Figure 5.**
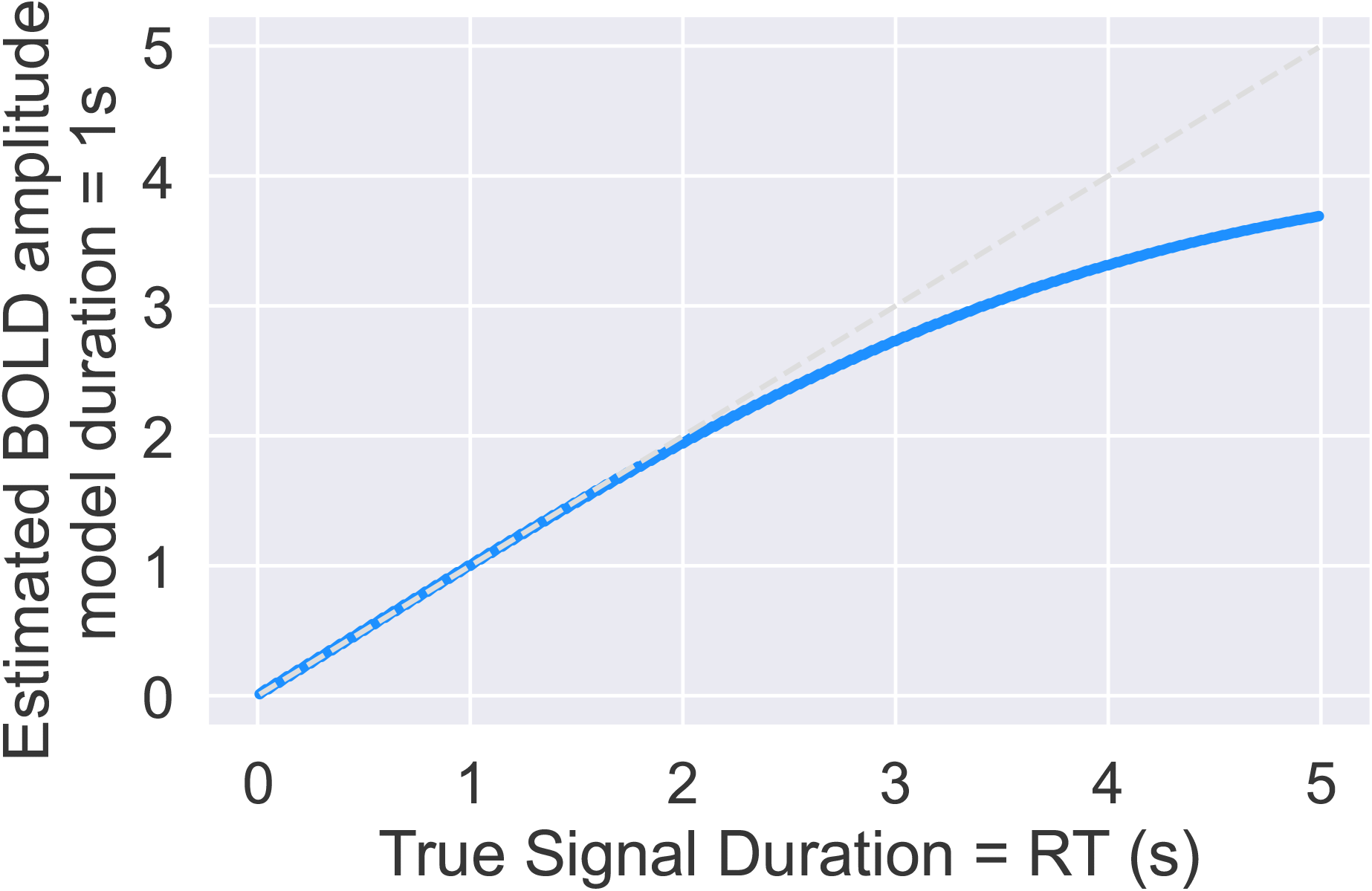
Relationship between trial RT and the trial-specific BOLD activation estimate when a constant duration of 1s regressor is assumed and signal scales with RT (blue). Gray dashed line is a line with a slope of 1 and intercept of 0. The true BOLD activation is 1, but the model estimates the BOLD activation to be 1 × ***RT*** for RTs < 2s.

The simulation results in Figure 6 show the relationship across all models and datatypes considered in this work. The ConstDurNoRT model produces correlations between contrast differences and RT differences when the signal scales with RT, as does the RTDur model when the signal does not scale with RT while the ConstDurRT model does not induce correlations for either signal type. In other words, although there is no true relationship between average subject RT difference and the fMRI contrast estimate, the ways in which these models cannot capture RT (ConsDurNoRT) or introduce RT information (RTDur) cause an RT effect at the group level, which may interfere with the correct interpretation of group level results (e.g., if group level variable of interest is related to RT). Notably, the data were simulated such that the variance in RT did not change with RT, whereas in real data the variance of RT often increases with its mean. The implication is the correlation estimated by our simulations is conservative. Even so, it is within the ballpark of the expected true correlations between brain and behavior measures (***Marek et al., 2022***).

**Figure 6.**
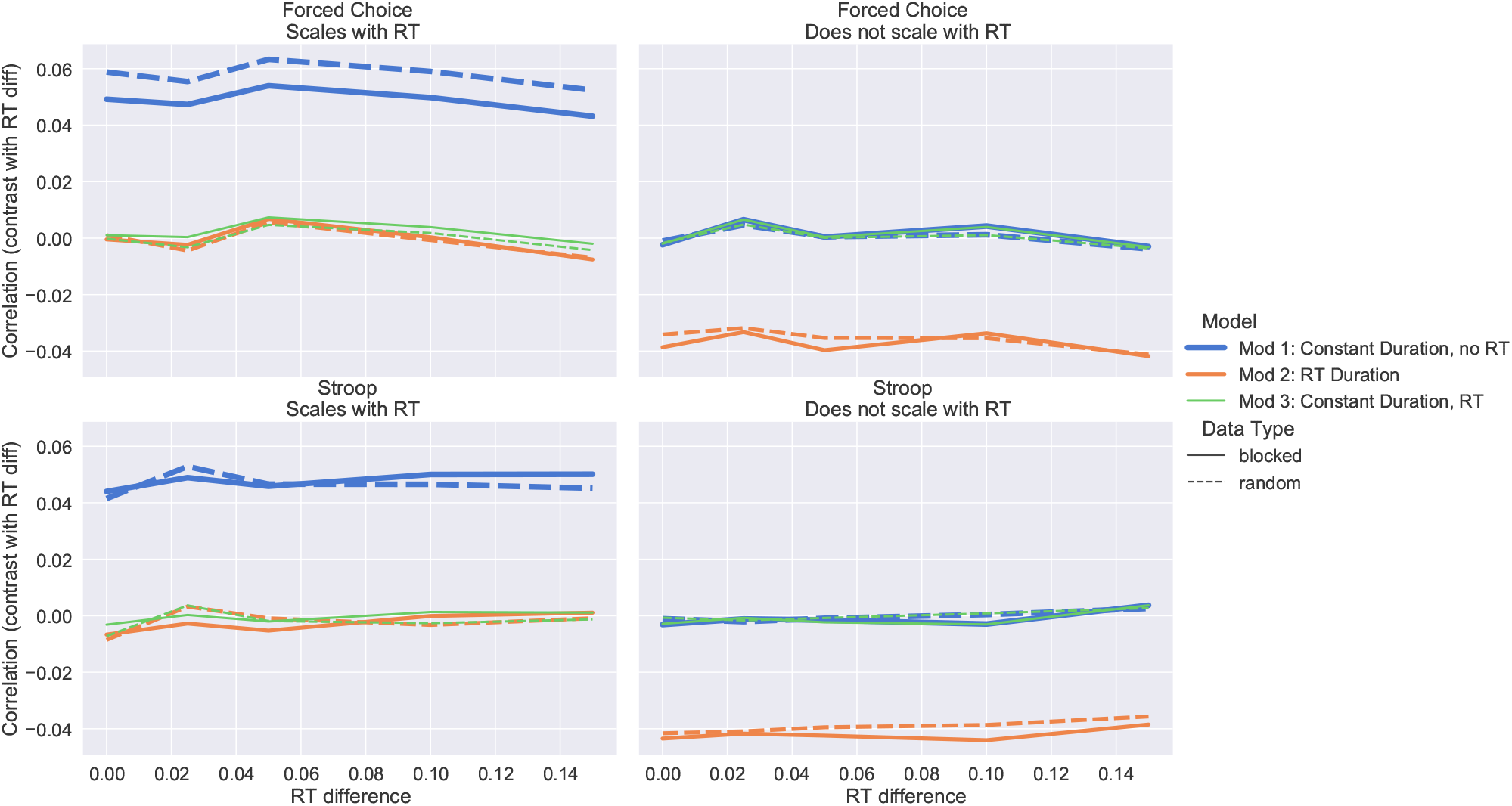
Correlation between the contrast difference and difference in average condition RT across subjects as a function of the average difference in RT between conditions. Since the correlation is driven by between-subject variability in the *difference* in RT, there is no requirement that RTs differ between tasks and the correlation is constant regardless of the RT difference.

Importantly in this scenario, the linear relationship at the group level is simply, *B,* the common activation for both conditions and all subjects. If the activation differs between conditions or across subjects the confound will potentially be more complex in the between-subject analysis and can even introduce new artifactual associations into a group analysis. A simple extension to illustrate how false associations can be introduced is to only relax the assumption that *B* is the same across subjects, but preserve the assumption that *B* is the same for both conditions. For example, assume age is equally related to both conditions through the relationship, *B* = *γ*_0_ + *γ*_1_*age* + *ϵ* and note that this does not introduce an association of age with the true activation difference. If the signal scales with RTs and the ConsDurNoRT model is used to estimate the condition difference, not only is an RT effect introduced to the group level analysis, but an artifactual age effect is also introduced. This can be seen in the following derivation that extends the earlier defined relationship with the new definition of *B*:

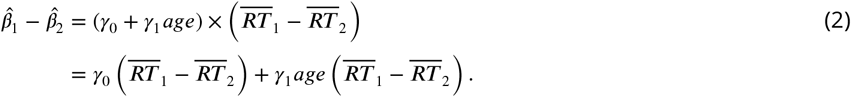

This result is concerning because there is not a true relationship of the condition difference with either the RT difference or age, but the use of ConsDurNoRT when the data scale with RT introduces an age association to the group level analysis through an interaction with the RT difference. The implication is if the RT difference is nonzero, on average, an age association will be present in the group level when there is not a relationship between age and the true activation difference. Adding RT difference as a confound regressor to the group model will not remedy these issues and should be avoided as it may inflate the significance of the false association with age. This calls into question the interpretation of between-subject analyses where the signal may scale with RT and ConsDurNoRT is used.

The relationship between RT and potential variables of interest will be different and more complex if, say, the RT difference is correlated with that variable or if there is a correlation between the true activation difference and the variable of interest. Just as in the example above, it is clear that adding an RT confound regressor to the group level model is not an adequate fix. The recommendation is to repeat the time series level analyses using ConsDurRT to avoid these issues. This is unsettling news for those using fMRI activation databases, since the ConsDurNoRT model is typically used to generate activation estimates.

### Widespread RT activation is not specific to task, revisited

Our real data analyses were modeled to included separate regressors for each condition as well as a single RT regressor to control for RT effects in our contrasts between conditions, similar to the ConsDurRT model. A total of 7 tasks, with sample sizes ranging from 86 to 94, were analyzed. The cognitive processes involved in these tasks include attention, temporal discounting, proactive control, reactive control, response inhibition, resisting distraction, and set shifting. Brief descriptions are given in Table 1 and more detailed summaries are provided in the Methods section. Comparatively, ***Yarkoni et al.*** (***2009***) used tasks including 3-back, decision making, emotion ratings and memory in sample sizes of 50, 102, 26, 35 and 39. Our seven tasks emphasize cognitive control to a greater extent and emotional processing and working memory to a lesser extent, compared to ***Yarkoni et al.*** (***2009***). The focus here is on the average RT-related effect across subjects. Notably, this effect estimate will be slightly diminished from a full RT effect, since it is adjusted for condition difference and so the interpretation would be the average within-condition RT effect. Group statistics maps were thresholded using the TFCE p-value (from FSL Randomise) less than 0.05 with 5000 permutations. The conjunction in Figure 7 shows voxels where the average RT-modulated effects were significant across all 7 tasks. Our maps are consistent with ***Yarkoni et al.*** (***2009***), but with a more spatially widespread effects, which may reflect the fact that our sample sizes were larger. In particular, the present comparison demonstrated substantially more signal in the lateral superior parietal cortex.

**Table 1.** fMRI task summaries

| Name | Description | N |
| --- | --- | --- |
| Attention Network Test (ANT) | Tests three aspects of attention or “attentional networks”: alerting, orienting, and executive control | 91 |
| Delay-Discounting Task (DDT) | Measure of temporal discounting, the tendency for people to prefer immediate monetary rewards over delayed rewards | 86 |
| Dot Pattern Expectancy (DPX) | Measure of individual differences in cognitive control including proactive and reactive control modes | 91 |
| Motor Selective Stop Signal | Measures the ability to engage response inhibition selectively to specific responses | 91 |
| Stop-Signal Task | Measure of response inhibition | 91 |
| Stroop | Measure of cognitive control perhaps including resisting distraction or attentional filtering | 94 |
| Cued Task-Switching Task (CTS) | Indexes the processes involved in reconfiguring the cognitive system to support a new task | 94 |

**Figure 7.**
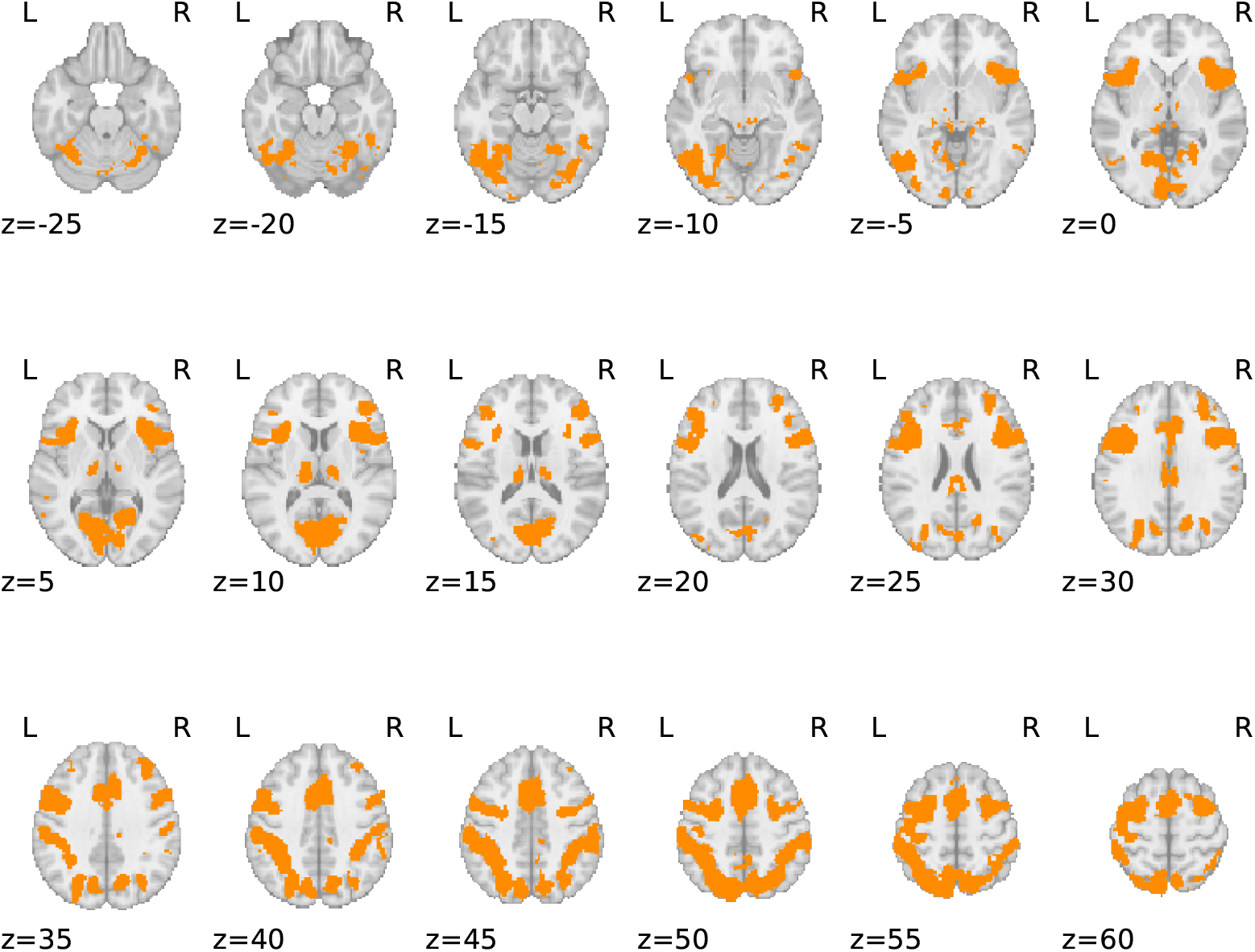
Conjunction of the average, within-condition RT effect across ANT, DDT, DPX, motor selective stop, stop signal, stroop, and CTS. On average, each analysis included around 91 subjects and maps were corrected for multiple comparisons using a TFCE p-value thresholded at 0.05 using 5000 permutations.

## Discussion

The problem of potential response time confounds for fMRI activation estimates has been discussed for more than a decade, but there has been little resulting change in how the community approaches the analysis and interpretation of fMRI contrasts when RT differences between task conditions are present. There are three takeaways from the present work. First, we propose a modeling approach that more effectively adjusts within-subject contrast estimates for response time differences, so that group average estimates of contrasts are adjusted for between-condition response time differences. Importantly this model does not remove the ability to also study RT-specific effects, if they are of interest. Second, this work highlights an important problem that has not been discussed previously: the presence of a between-subject analysis confound of the average RT differences and the potential to introduce artificial associations with variables of interest at the group level. Finally, we replicate previous work showing that RT-related effects are not task specific (***Yarkoni et al., 2009***).

This work presents a model that can adapt to data whether or not the signal scales with RT, without losing performance. By adding an RT modulated regressor to the most commonly used model that only contains condition-specific regressors, the ConstDurRT model reduces RT-driven type I errors in average condition comparison effects without a reduction in power. The commonly used ConstDurNoRT model assumes the signal does not scale with RT and the RTDur model assumes the signal must scale with RT, hence both models fail to control error rates when these model assumptions are violated (Figure 3).

We have also uncovered that subject-specific differences in average RT represent an important group-level confound. This confound is only present if the time series model follows the ConsDurNoRT or RTDur approaches, whereas the ConsDurRT model produces condition difference effects that are free of this confound. Notably, the average difference in RT across subjects does not impact the strength of the correlation, since the correlation is driven by the variability in RT differences across subjects. Thus, even if the RT difference is 0, the first level models should include a between-trial RT adjustment. We have also shown that when the signal scales with RT and the ConsDurNoRT model is used, other false associations can be introduced at the group level. Specifically we have shown that when there is a common association between a variable of interest (e.g., age) and each condition, separately, but no association of this variable with the true condition difference, the ConsDurNoRT condition difference estimates can have false associations with this variable. This is an especially concerning result for users of neuroimaging databases of condition difference estimates, since ConsDurNoRT is typically used.

Last, our updated RT-effect conjunction analysis across 7 tasks tapping into different mental processes show widespread shared activation in the so-called “task-positive” network, replicating the previous results of ***Yarkoni et al.*** (***2009***). This highlights the generality of the RT effect across tasks, and motivates the need to model these effects across all tasks.

### The paradox that RT is the effect of interest in fMRI studies

Our recommendation here is to focus separately on RT-based effects and condition differences, adjusted for RT. There can be resistance to this idea, since RT is the measure of interest in behavioral studies and removing RT effects from condition differences is argued to be “throwing the baby out with the bathwater”. This argument is paradoxical since if RT effects are the effects of interest, then why not study the RT effects directly? Studying unadjusted condition differences does not directly reflect RT effects and may reflect differences that are completely unrelated to RTs. In fact the condition difference effect, when unadjusted for RTs, may be driven by any of the underlying models shown in Figure 8 and this fact should be made clear when presenting any results focused on averages of condition differences estimated with the ConsDurNoRT model. If the RT based effect is the effect of interest, it should be studied directly using a model that mirrors the underlying theoretical model of RTs relationship with the construct of interest to maximize power. A significant finding of an RT-based effect is just the beginning and further work is necessary to establish whether the RT-based effect reflects time on task or an amplitude-driven effect, as was argued for the Stroop task by ***Yeung et al.*** (***2011***). If this important extra work is not done, then conclusions for unadjusted RT effects must be presented acknowledging the limitations in their interpretation.

**Figure 8.**
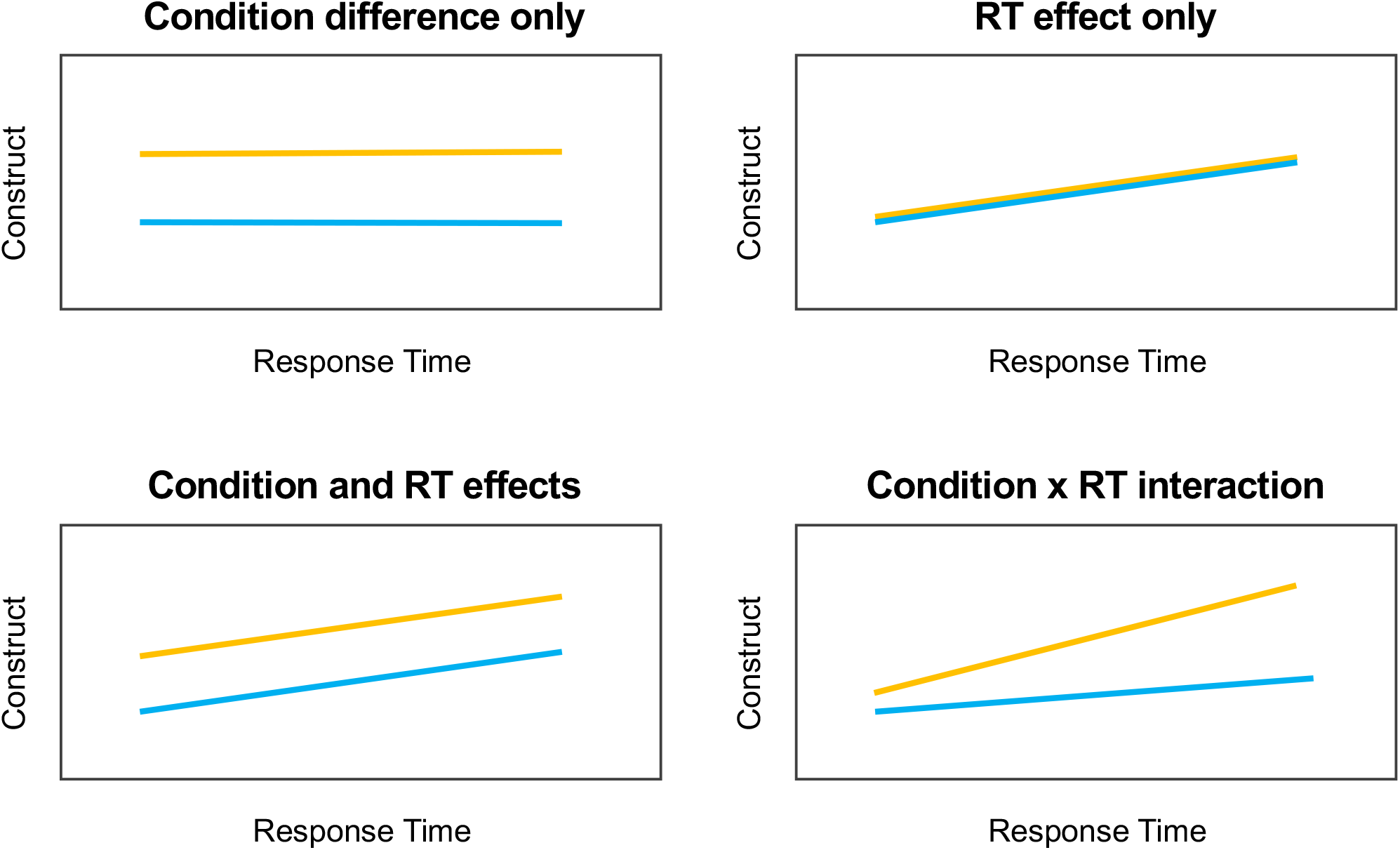
The ConsDurNoRT model assumes the underlying relationship between the construct of interest follows the “Condition difference only model. Since the condition difference model can yield significant results when *any* of these models is the true underlying model, only nonspecific interpretations can be made from the ConsDurNoRT model results. To strongly link brain activation to a specific construct of interest, the fMRI model should mirror the underlying behavioral theoretical model.

Overall our recommendation of focusing on RT-based effects and condition differences, adjusted for RTs, separating the two effects so they can be more accurately interpreted, ultimately improving our understanding of the brain.

### Modeling considerations

#### Should the RT modulation values be centered?

We did not center RT in our models, as it would not have any impact on the condition difference estimate since trials in both conditions involve RTs and the model implies the same condition difference effect occurs for all RTs. A common practice is to center by the mean RT for that subject and run of data, but this can introduce RT information into some contrast estimates. For example, if RT is centered, the interpretation of a condition versus baseline effect is specifically for that subject/run’s mean RT. In this case there is an RT confound introduced at the group level since each subject’s activation reflects their own RT. The same will occur for any contrast where one condition involves RTs and another does not. For example, in the stop signal task the go trials have a response time whereas successful stop trials do not. Therefore if RT is centered within-subject and run, the go versus successful stop contrast estimate corresponds to the magnitude of the effect for mean RT of that subject/run, such that a correlation between this contrast and the average go RT will leak into the group level analysis. A summary of contrast types that are impacted by centering RT is given in Table 2.

**Table 2.** Whether or not centering of the RT modulated regressor is necessary when using ConsDurRT to study adjusted condition differences. When centering is required do not use the mean RT for the run, but use the same centering value for all subjects and runs to prevent incorporating an RT confound in between-subject analyses.

| Contrast type | Example<br>(stop signal task) | Interpretation without<br>centering RT | RT centering<br>necessary with<br>ConsDurRT model? |
| --- | --- | --- | --- |
| Condition (RTs) vs.<br>baseline | Go vs.<br>baseline | Go activation when RT<br>is 0 | Yes |
| Condition (RTs) vs.<br>Condition (no RTs) | Go vs.<br>Successful Stop | Condition difference<br>when Go RT is 0 | Yes |
| Condition (RTs) vs.<br>Condition (RTs) | Go vs.<br>Unsuccessful Stop | Condition difference | No |

To avoid this issue simply center by the same value for all subjects and runs. This value can either be the mean RT across *all* subjects and runs or a value that is roughly what one would expect the average RT to be for that task. In this case the RT confound at the group level should not be present. If all condition comparisons involve conditions with RT effects (as is most often the case for contrasts of interest in cognitive fMRI studies), then centering RT in the modulated regressor will have no effect on the contrast estimates of interest.

#### Why an RT modulation is used instead of RT duration

It may seem counterintuitive that we model RT through a parametrically modulated regressor in the ConsDurRT model, instead of a single RT duration style regressor. This is because the RT centering, described in the previous section, is a special type of orthogonalization that is not possible to carry out directly with the RT duration style regressors in most fMRI software packages in a straightforward way. Efforts to do so will typically reflect an unintended orthogonalization that removes condition difference information from the RT regressor, which should be avoided. Even if orthogonalization could be done properly, it will cause the condition-based activation to reflect subject-specific RT averages in the same way as mean centering was described to have issues in the previous section. Generally the RT modulated regressor is a very close approximation to the RT duration regressor, so it is an excellent substitute with more flexibility so we can properly adjust it to improve the interpretation of condition difference contrasts without introducing a new between-subject RT confound.

#### Avoiding common pitfalls when adding RT to a time series model

When adding RT to the time series model, there are some common mistakes that should be avoided. Most of the problem relates to the intuition that collinearity between regressors is problematic and thus that mean centering or orthogonalization is always necessary. Once the data have been collected, collinearity with regressors of interest often cannot be resolved, but may have been preventable with a different study design. If the RT regressor is highly collinear with the task contrast, this should not be altered by orthogonalization or centering RT within task, as that is in conflict with the motivation for adding RT in the first place: controlling condition differences for RT differences or time on task effects. If RT is mean centered within-condition, then it is misleading and incorrect to claim the condition effects have been adjusted for RT differences. If RT is split into separate regressors by condition, this model implies an interaction effect between condition and task is suspected. In this case, if the interaction is significant, then that is the contrast to be studied in detail as it indicates the magnitude of the condition difference varies by RT. One should not use an interaction model to study main condition effects, even if the interaction is not found to be significant, which is discussed in the next section. Our recommendation is if there is not an expected interaction between different conditions and RT, then a single RT regressor should be used.

#### Condition by RT interaction models

If the underlying theory about the relationship between the psychological measure of interest and RT implies a condition by RT interaction, then an interaction model should be used in the fMRI analysis. Specifically the single RT regressor from ConsDurRT would be split by condition and no RT centering should be applied. In this case the effect of interest is the difference in the parameters corresponding to the RT modulated regressors for each condition. If the interaction is not found to be significant, we discourage using the interaction model to then study the main condition effects, as recommended in ***Carp et al.*** (***2010***). Although finding the interaction is not significant means the estimated slopes between RT and BOLD activation are not statistically different different between conditions, the estimated slopes will be different and this adds noise into the condition difference estimate, which reduces power. Instead, to study main condition differences when the interaction is not found to be significant, we recommend simplifying to the ConsDurRT model. A group level power analysis (Figure S2) indicates that the additional noise introduced when using the interaction model results in a loss in power as high as 9.5% compared to the ConsDurRT model (Figure S2). Of course, to avoid inflated error rates, p-value thresholds should use a correction for multiple comparisons (e.g., Bonferroni correction) to correct for testing both the interaction and then the main condition effect if the interaction is not found to be significant.

### Limitations of this work

This work consists of real data analyses as well as simulated data analyses. Simulations are required in cases where we need to know the ground truth and link the theoretical problems with how these problems might surface in real data analyses (e.g., how strong the results are and whether they persist at the group level). As such, the simulations require specifying a large number of parameters including the RT distribution for each condition, effect size for each condition, stimulus length, ISI, within-subject variance and between-subject variance. We specifically focus on the RT distribution used in previous work (***Grinband et al., 2008***) to broaden those conclusions to more models and add an RT distribution based on our own Stroop data, which is quite different. In an effort to set the rest of the parameters to realistic values we focused on the size of the within-subject condition effect, aiming for correlations 0.07-0.08, ratio of total variance to within-subject variance, 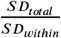, ranged between 2-3 and the Cohen’s D for the average of task versus baseline across subjects was approximately 0.85. Higher between-subject variance (lower 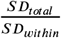) would yield smaller time on task effects. Additionally we only shifted the mean parameters of the Exponential Gaussian distributions used to define the RT differences between conditions, whereas our analysis of our own Stroop data showed the variance increased with average within-subject RT. This was done intentionally to isolate how mean RT differences impact different analyses while holding the RT variance constant. This increased variance in RT as RT increases would only strengthen the between-subject correlation between the condition contrast and RT difference at the group level. Even though these limitations exist, we believe our results to be accurate representations of the RT effect based on consistencies with other studies focusing on the RT effect (***Yarkoni et al., 2009***; ***Brown, 2011***; ***Grinband et al., 2011b***).

## Methods

### Models considered

#### Data generation and modeling

The interstimulus interval (ISI) was sampled from a Uniform distribution and RT was sampled from an ex-Gaussian distribution. For RT, a subject specific *μ_sub_* was obtained by sampling an ex-Gaussian with parameters *μ_rl_, σ_rl_* and 1/*λ_rl_* and subtracting the sampled value by 1/*Λ_rl_.* The subject-specific RTs were then sampled from an ex-Gaussian distribution with *μ_sub_, σ_rt_* and 1/2_*rt*_. When RT differed between conditions, each Condition 1’s RT mean was *μ_sub_*-Δ*RT*/2 and Condition 2’s was *μ_sub_* + Δ*RT*/2, where Δ*RT* was the RT difference. Values of *μ_n_, σ_n_* and *λ_r1_* were based on our Stroop data and the Forced Choice Task in ***Grinband et al.*** (***2008***). In both cases distributions were fit to subject-specific data and then parameters were averaged over subjects. The Forced Choice RT distribution was defined by a Gamma distribution with shape parameter = 1.7, beta = 0.49. Sampling from this distribution and fitting an ex-Gaussian to that sample resulted in ex-Gaussian parameters of *μ_rt_* = 638, *σ_rt_* = 103, and 1/*λ_rl_* = 699 (mean = 1337, sd = 706.5). The Stroop data had faster RTs with less variability, with ex-Gaussian parameters of *μ_rt_* = 530, = 77, and 1/2_*rt*_ = 160 (mean = 690, sd = 177.5). The distribution functions from Python’s Scipy module were used to simulate and estimate the distribution parameters. Trials were either randomly presented conditions or blocked conditions, where 4 trials of the same condition were presented in a row.

Simulated data that scaled with RT were created with the convolved RT duration regressors (RTDur) and data that did not scale with RT used the constant duration regressors (ConsDurNoRT). The BOLD activation sizes for the *i^th^* subject for each condition, *β*_*i*,1_ and *β*_*i*,2_, were sampled from a Gaussian distribution, 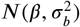, where *β* is the true activation magnitude and 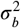 is the between-subject variance. The time series data for the *i^th^* subject, of length *T*, was created according to

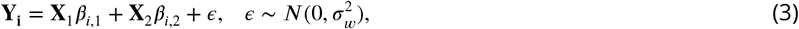

where **X**_1_ and **X**_2_ are either the Model 1 or 2 regressors (*T* × 1) and 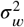 is the within-subject variance.

In an effort to choose realistic values for *β*_1_, *β*_2_, 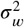 and 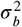, we considered the first level effect size (converting the true *β_i_* to a correlation), second level effect size for a 1-sample t-test (Cohen’s D) as well as the ratio of the total mixed effects variance to the within-subject variance. Following the definitions of parameters as given in the model above, the total mixed effects variance for a first level contrast of parameter estimates is

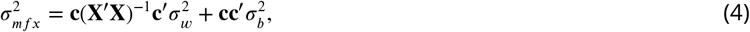

where **X** and **c** are the first level design matrix (based on models in Figure 2) and contrast of interest (***Mumford and Nichols, 2006***). The contrast of interest for each model corresponded to condition 2 > condition 1 (**c** = [-1,1] for the 2 regressor models and **c** = [-1,1,0] for the three regressor model). The ratio of total standard deviation (SD) to within-subject SD is defined by

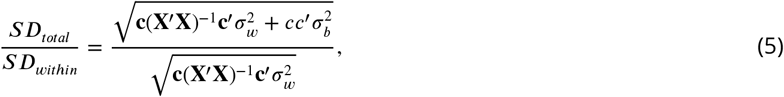

Our within-subject effect size for condition versus baseline was between 0.07-0.08 (correlation), ratio of total variance to within-subject variance, 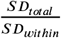, ranged between 2-3 and the Cohen’s D for the average of task versus baseline across subjects was approximately 0.85.

Each run contained 40 trials of each condition and a time resolution (TR) of 1s. Time course length varied, as it was set to extend 50s past the last stimulus offset. Group analyses included 100 subjects. A total of 1000 data sets were simulated to calculate power and error rates.

Regressors were constructed by convolving boxcar functions with a Double Gamma hemodynamic response function (HRF) using the compute_regressor function from the Nilearn module in Python.

Least squares regression was used to estimate the models described in Figure 2 at the first level including a set of cosine basis functions (0.1 Hz cutoff) for high-pass filtering generated with the cosine_drift function from Nilearn in Python. At the group level, 1-sample t-tests were used to assess type I error and power. A correlation of the average difference in RT between conditions and the fMRI contrast (condition 2 vs condition 1) was estimated for each group analysis.

Since RT and ISI values are random, the contribution of the design matrix, *X,* to the overall variance varies between samples (Equation 4) and the true effect size was variable. Therefore, to calculate the first level true effect size 100 data sets were simulated and the partial correlation coefficient for one condition, controlling for the other condition and cosine basis set, was estimated and then averaged over the 100 data sets to serve as the true within-subject effect. The variance ratio, *SD_total_/SD_within_*, was estimated by simulating 100 design matrices. Cohen’s D estimates were based on 5000 simulated within-subject model estimates for the task versus baseline contrast.

#### Real data analysis

A total of 110 subjects completed each of the following fMRI tasks: Stroop (***Stroop, 1935***), Attention Network Test (ANT, ***Fan et al.*** (***2002***)), Dot Pattern Expectancy task (DPX, ***MacDonald et al.*** (***2005***)), Delayed-Discounting task (DDT, ***Kirby*** (***2009***)), cued task-switching task (CTS, ***Logan and Bundesen*** (***2003***)), stop signal task (***Logan and Cowan, 1984***) and a motor selective stop signal task (***DeJong et al., 1995***). Brief summaries are provided in Table 1 and more detailed descriptions are provided in the Supplementary materials. Data were acquired using single-echo multi-band EPI. The following parameters were used for data acquisition: TR = 680ms, multiband factor = 8, echo time = 30 ms, flip angle = 53 degrees, field of view = 220 mm, 2.2 × 2.2 × 2.2 isotropic voxels with 64 slices.

Data were preprocessed in Python using fmriprep 20.2.0 (***Esteban et al., 2019***). First, a reference volume and its skullstripped version were generated using a custom methodology of fMRIPrep. A B0-nonuniformity map (or fieldmap) was directly measured with an MRI scheme designed with that purpose (typically, a spiral pulse sequence). The fieldmap was then co-registered to the target EPI (echo-planar imaging) reference run and converted to a displacements field map (amenable to registration tools such as ANTs) with FSL’s fugue and other SDCflows tools. Based on the estimated susceptibility distortion, a corrected EPI (echo-planar imaging) reference was calculated for a more accurate co-registration with the anatomical reference. The BOLD reference was then co-registered to the T1w reference using bbregister (FreeSurfer) which implements boundary-based registration (***Greve and Fischl, 2009***). Co-registration was configured with six degrees of freedom. Head-motion parameters with respect to the BOLD reference (transformation matrices, and six corresponding rotation and translation parameters) are estimated before any spatiotemporal filtering using mcflirt(FSL 5.0.9, ***Jenkinson et al.*** (***2002***)). BOLD runs were slice-time corrected using 3dTshift from AFNI 20160207 (***Cox and S*** (***1997***), `RRID:SCR_005927`). The BOLD time-series (including slice-timing correction when applied) were resampled onto their original, native space by applying a single, composite transform to correct for head-motion and susceptibility distortions. These resampled BOLD time-series will be referred to as preprocessed BOLD in original space, or just preprocessed BOLD. The BOLD time-series were resampled into standard space, generating a preprocessed BOLD run in MNI152NLin2009cAsym space. First, a reference volume and its skull-stripped version were generated using a custom methodology of fMRIPrep. Automatic removal of motion artifacts using independent component analysis (ICA-AROMA, ***Pruim et al.*** (***2015***)) was performed on the preprocessed BOLD on MNI space time-series after removal of non-steady state volumes and spatial smoothing with an isotropic, Gaussian kernel of 6mm FWHM (full-width half-maximum). Corresponding “non-aggresively” denoised runs were produced after such smoothing. These data were used in our time series analysis models.

Data were analyzed using FirstLevelModel from nilearn in Python (***Abraham et al., 2014***). A double gamma HRF was used for convolution and an AR(1) model addressed temporal auto correlation. Regressors were included for each condition, versus baseline, as well as a single RT modulated regressor, similar to the simulation analysis model ConsDurRT. The RT modulated regressor included the uncentered RT values. The contrast of the RT modulated regressor was the contrast of interest in our models and represents the average relationship between BOLD activation and RT within condition, since condition specific regressors were also included. Nuisance regressors in the time series analysis included the following from the fmriprep output: cosine basis functions (corresponding to a highpass filter cutoff of 128s) and the average time courses for the CSF and WM as estimated by fmriprep.

Subjects were excluded within-task for the following general reasons: missing 1 or more files required to analyze the data, having more than 20% high motion time points (measured by Framewise Displacement > .5 or SD of DVARS > 1.2), having more than 45% missing responses, a subjective poor performance rating assessing high choice and/or omission error rates in at least one condition of the task, and when subjects omitted most of their responses towards the end of the task scan. Specific exclusion for the stop signal tasks are less than 25% successes for stop trials or more than a 75% successful stop rate. For the Delay-Discounting tasks, subjects were excluded if they made the same choice on all trials. Last, if there were exclusions on more than half the tasks for a subject, that subject was completely excluded. For Stroop 9 subjects had missing data files, 1 subject had more than 45% missing responses, 2 subjects had >20% high motion volumes, 1 subject had exclusions on more than half the tasks, 1 subject had exclusions

Group models were estimated using Randomise (***Smith and Nichols, 2009***) and included either a single column of 1s (group mean) or a column of 1s along with the difference in mean RTs. Statistics maps were thresholded, controlling for family-wise error rate, using the Randomise TFCE statistic below 0.05, based on 5000 permutations. Two sided hypotheses were studied using an F-contrast. A conjunction map was constructed by taking the overlap of the thresholded, binarized map for each of the 7 tasks (***Nichols et al., 2005***).

## Acknowledgements

We would like to thank Daniel Weissman for discussions that helped in the writing of this manuscript.

## Supplement

### Error rate when ISI ranges between 3-6s

**Figure S1.**
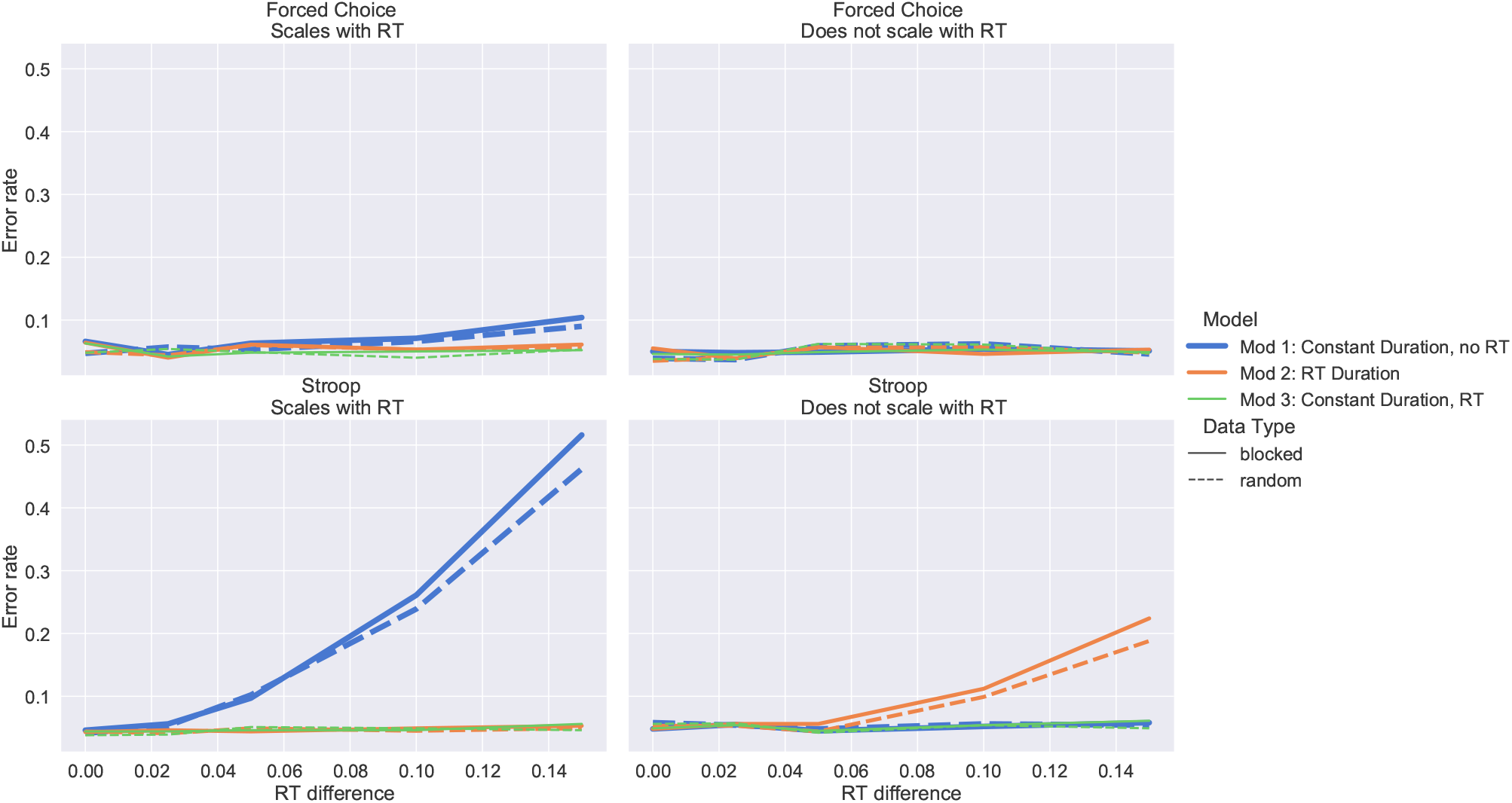
Type I error as RT difference between conditions increases. This illustrates that results are similar to when the ISI ranged between 2-4s (result in main manuscript). The Forced Choice Task RT distribution was used in the top panels, while Stroop RT distribution was used in the bottom panels, both with an ISI between 3-6s was used and inference of interest was the 1-sample t-test of the condition effect with 100 subjects. 2500 simulations were used to calculate the error rate.

### Power differences when studying condition differences after testing for interaction

Here we study the power for a condition effect after testing for a potential condition by RT interaction. The interaction model contained two condition regressors and two RT regressors, split by condition. RTs were centered by the theoretical RT based on the distribution used to simulate RTs. Although the slopes of the interaction model were not found to significantly differ, their magnitudes will not be exactly equal and this introduces variance into the condition difference estimate from this model. This is reflected in the reduced power compared to the ConsDurRT model (Figure S2).

**Figure S2.**
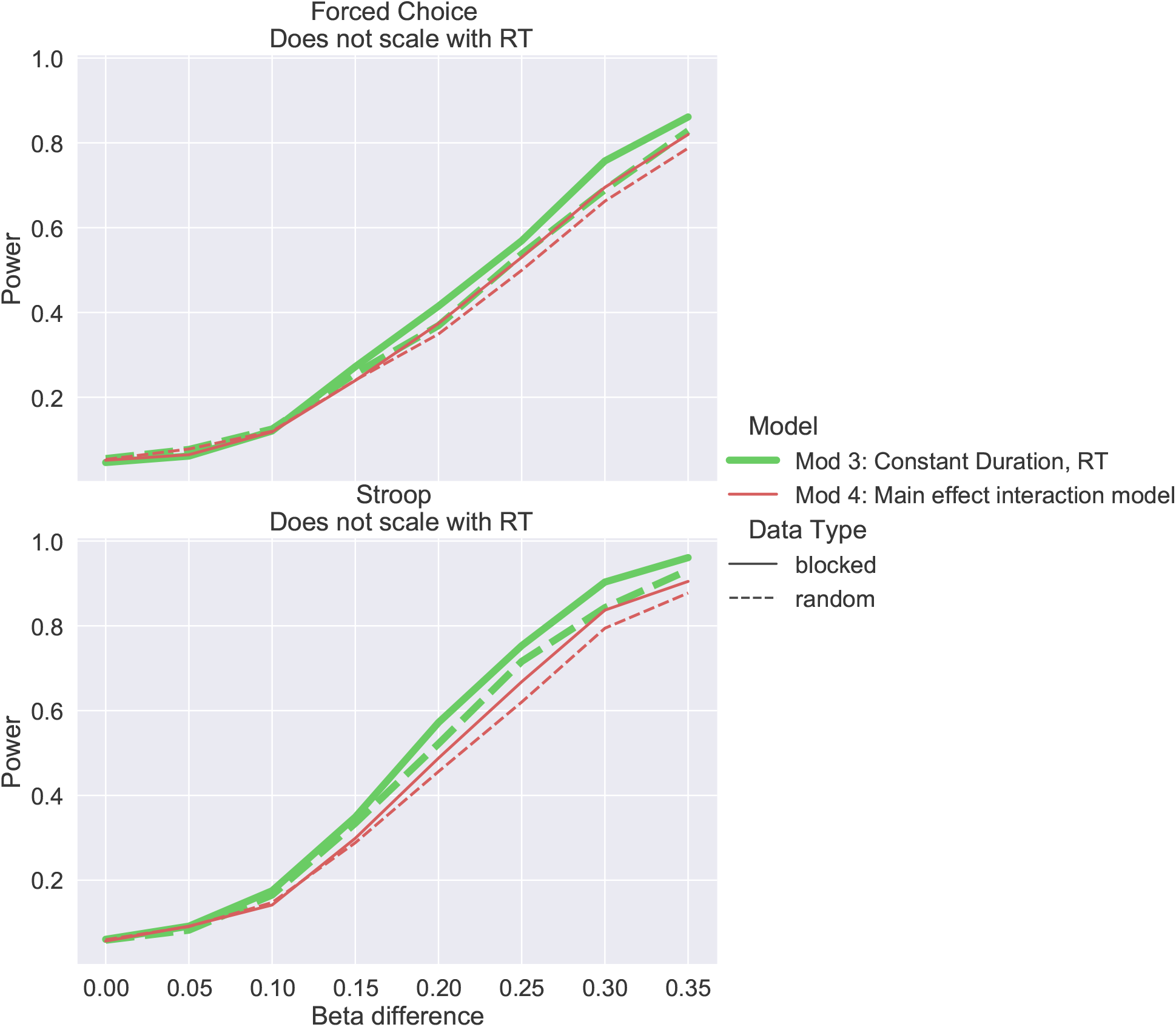
Power for testing main condition difference effect after an interaction was found to be not significant. Power is lost if the main effect of condition is studied in the interaction model, directly, due to the estimated sloped not being equal, which introduces variability into the estimate. Power is improved if the condition difference is then studied using the ConsDurRT model. Note, the p-value cutoff for testing the interaction and main effect of condition were set to .025 to preserve the overall error rate at 5%.

### Details about tasks involved in real data analysis

The Attention Network Test (ANT) is a task designed to test three attentional networks: (1) alerting, (2) orienting, and (3) executive control. The ANT combines attentional and spatial cues with a flanker task (a central imperative stimulus is flanked by distractors that can indicate the same or opposite response to the imperative stimulus). On each trial a spatial cue is presented, followed by an array of five arrows presented at either the top or the bottom of the computer screen. The subject must indicate the direction of the central arrow in the array of five. The cue that precedes the arrows can be non-existent, a center cue, a double cue (one presented at each of the two possible target locations), or a spatial cue that deterministically indicates the upcoming target location. Each network is assessed via reaction times (RTs). The alerting network contrasts performance with and without cues, the orienting network contrasts performance on the task with or without a reliable spatial cue, and executive control (conflict) is measured by assessing interference from flankers.

The Dot Pattern Expectancy (DPX) task measures individual differences in cognitive control. Participants are presented with a cue made up of dots. This cue can be a valid cue – referred to as A (e.g.,“:”) – or an invalid cue – referred to as B (e.g.,“..”). Next a probe is presented, also made up of a simple dot formation. This probe can be valid (X) or invalid (Y). Participants are instructed to respond to valid probe and cue combinations (targets – AX combinations) with a key press (e.g., “x”) and all others (non-targets) with a different key press (e.g., “m”).

The Delay-Discounting Task (DDT) is a measure of temporal discounting, the tendency for people to prefer smaller, immediate monetary rewards over larger, delayed rewards. Participants complete a series of 27 questions that each require choosing between a smaller, immediate reward (e.g., $25 today) versus a larger, later reward (e.g., $35 in 25 days). The 27 items are divided into three groups according to the size of the larger amount (small, medium, or large). Modeling techniques are used to fit the function that relates time to discounting. The main dependent measure of interest is the steepness of the discounting curve such that a more steeply declining curve represents a tendency to devalue rewards as they become more temporally remote.

The cued task-switching task indexes the control processes involved in reconfiguring the cognitive system to support a new stimulus-response mapping. In this task, subjects are presented with a task cue followed by a colored number (between 1-4 or 6-9). The cue indicates whether to respond based on parity (odd/even), magnitude (greater/less than 5), or color (orange/blue). Trials can present the same cue and task, or can switch the cue or the task. Responses are slower and less accurate when the cue or task differs across trials (i.e., a switch) compared to when the current cue or task remains the same (i.e., a repeat).

The Stop-Signal Task is designed to measure motor response inhibition, one aspect of cognitive control. On each trial of this task participants are instructed to make a speeded response to an imperative “go” stimulus except on a subset of trials when an additional “stop signal” occurs, in which case participants are instructed that they should make no response. The Independent Race Model describes performance in the Stop-Signal Task as a race between a go process that begins when the go stimulus occurs and a stop process that begins when the stop signal occurs. According to this model, whichever independent process reaches completion first determines the resulting behavior; earlier completion of the go process results in an overt response (i.e., stop-failure), whereas earlier completion of the stop process results in successful inhibition. The main dependent measure, stop-signal reaction time (SSRT), can be computed such that lower SSRT indicates greater response inhibition. One variant of the task measures proactive slowing, the tendency for participants to respond more slowly in anticipation of a potential stopping signal. This variant often uses multiple probabilities of a stop signal (e.g., 20% and 40%) to manipulate participants’ expectancies about the likelihood of a stop signal occurring. The extent of slowing in the higher compared to the lower stop probability conditions is an index of proactive slowing/control.

The motor selective stop-signal task measures the ability to engage response inhibition selectively to specific responses. In this task, cues are presented to elicit motor responses (e.g., right hand responses, left hand responses). A stop-signal is presented on some trials, and subjects must stop if certain responses are required on that trial (e.g., right hand responses) but not others (e.g., left hand responses) if a signal occurs. In contrast to a simple stop-signal task in which all actions are stopped when a stop-signal is presented, this task aims to be more like stopping in “the real world” in that certain motor actions must be stopped (e.g., stop pressing the accelerator at a red light) but others should proceed (e.g., steering the car and/or conversing with a passenger). Commonly, stop-signal reaction time (SSRT), the main dependent measure for response inhibition in stopping tasks, is prolonged in the motor selective stopping task when compared to the more canonical simple stopping task. This prolongation of SSRT is taken as evidence of the cost of engaging inhibition that is selective to specific effectors or responses.

The Stroop task is a seminal measure of cognitive control. Successful performance of the task requires the ability to overcome automatic tendencies to respond in accordance with current goals. On each trial of the task, a color word (e.g., “red”, “blue”) is presented in one of multiple ink colors (e.g., blue, red). Participants are instructed to respond based upon the ink color of the word, not the identity of the word itself. When the color and the word are congruent (e.g., “red” in red ink), the natural tendency to read the word facilitates performance, resulting in fast and accurate responding. When the color and the word are incongruent (e.g., “red” in blue ink), the strong, natural tendency to read must be overcome to respond to the ink color. The main dependent measure in the Stroop task is the “Stroop Effect”, which is the degree of slowing and the reduction in accuracy for incongruent relative to congruent trials.

### Exclusion information by task for real data analysis

**Table S1.**
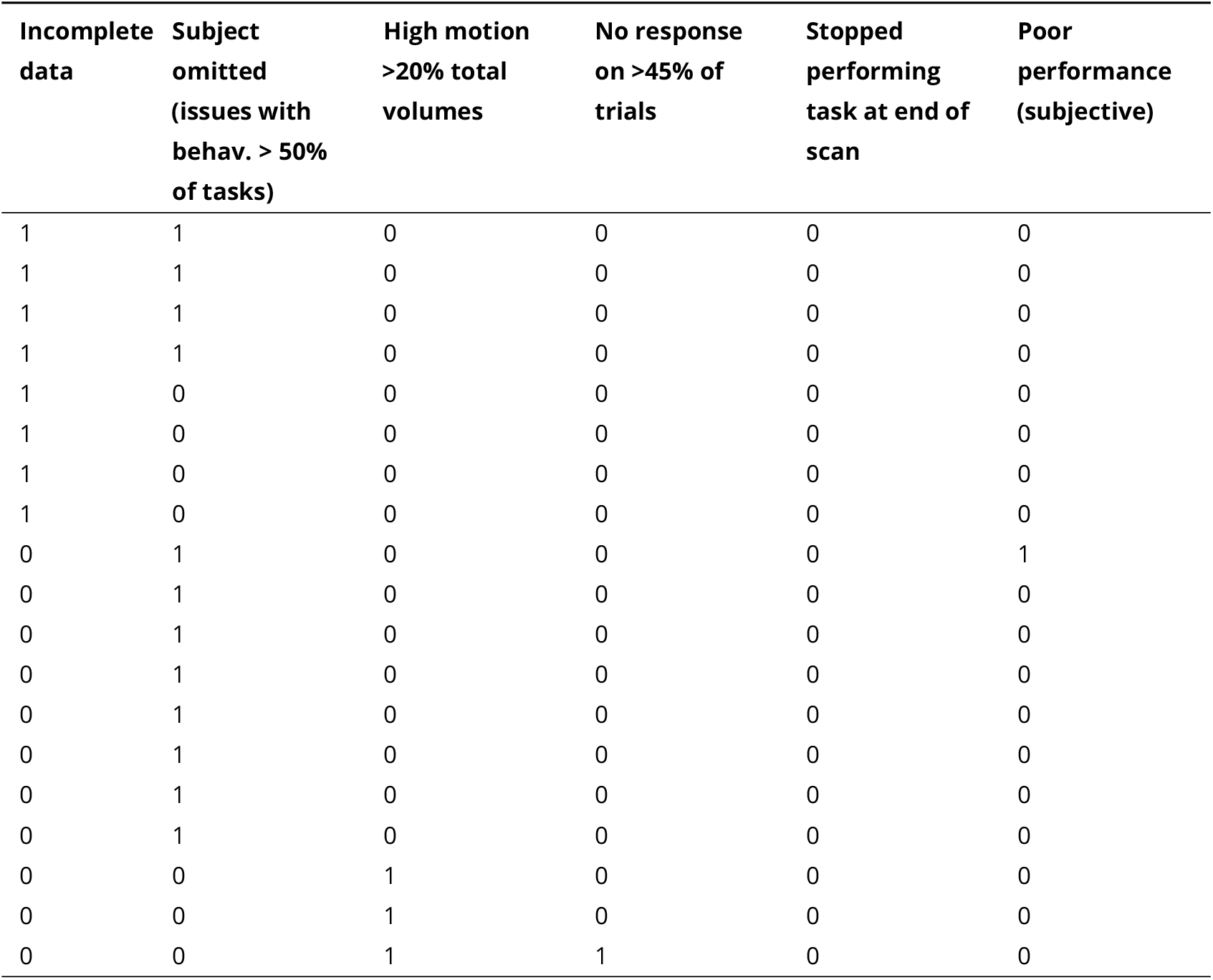
Exclusion information for Attention Network task.

**Table S2.**
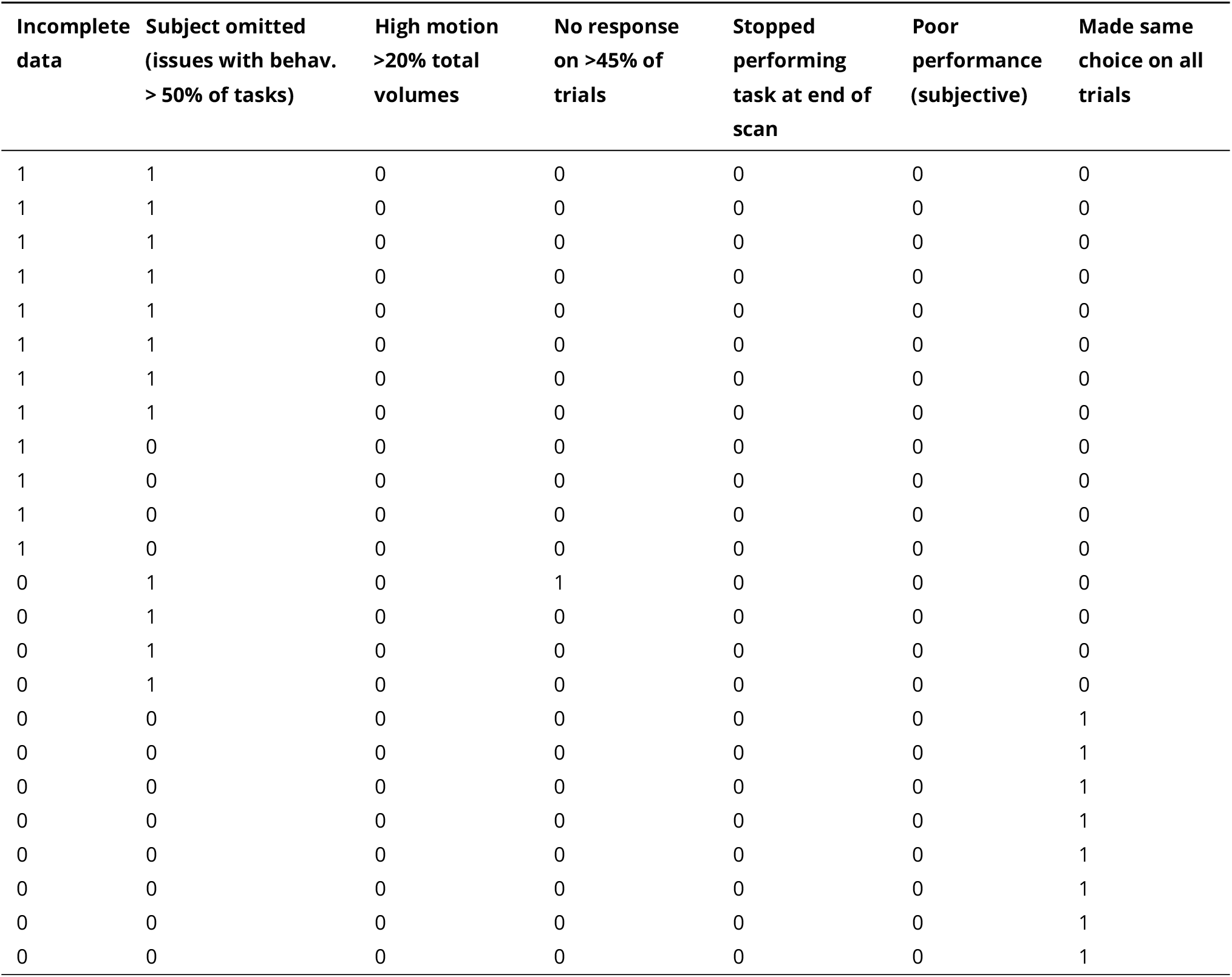
Exclusion information for Delay-Discount task.

**Table S3.**
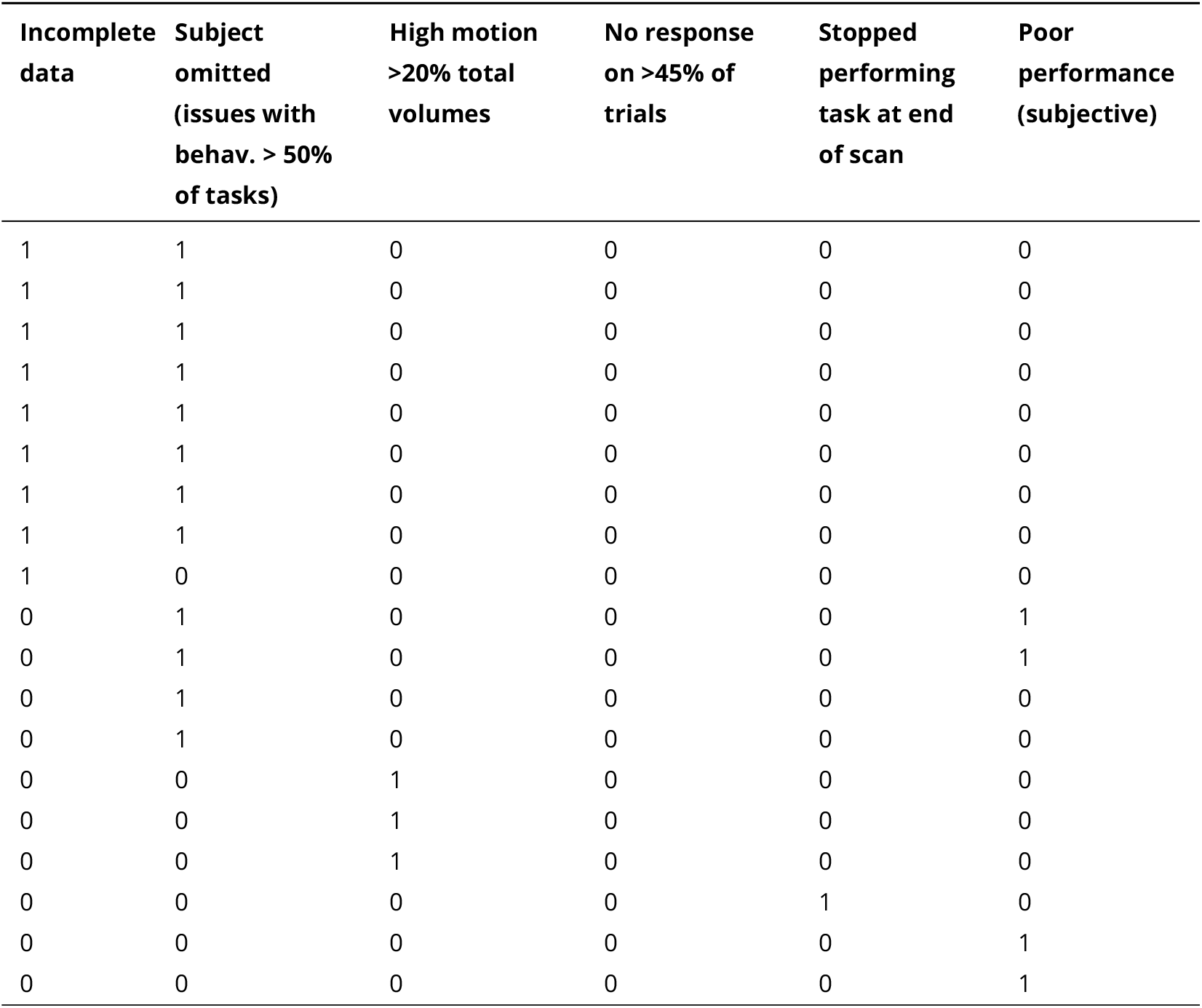
Exclusion information for Dot Pattern Expectancy task.

**Table S4.**
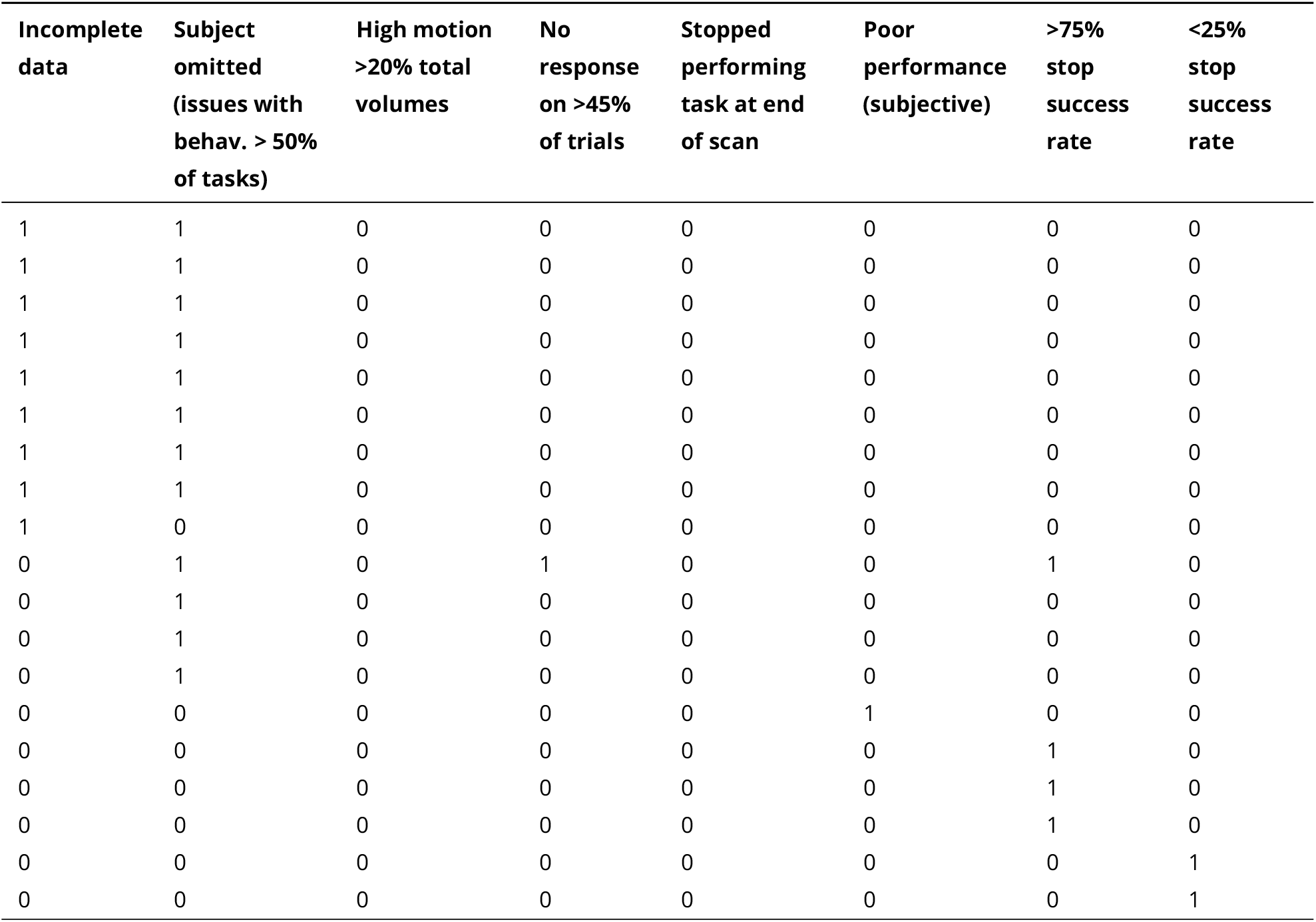
Exclusion information for Motor Selective Stop Signal task.

**Table S5.** Exclusion information for Stop Signal task.

| **Incomplete data** | **Subject omitted (issues with behav. > 50% of tasks)** | **High motion >20% total volumes** | **No response on >45% of trials** | **Stopped performing task at end of scan** | **Poor performance (subjective)** | **>75% stop success rate** | **<25% stop success rate** |
| --- | --- | --- | --- | --- | --- | --- | --- |
| 1 | 1 | 0 | 0 | 0 | 0 | 0 | 0 |
| 1 | 1 | 0 | 0 | 0 | 0 | 0 | 0 |
| 1 | 1 | 0 | 0 | 0 | 0 | 0 | 0 |
| 1 | 1 | 0 | 0 | 0 | 0 | 0 | 0 |
| 1 | 1 | 0 | 0 | 0 | 0 | 0 | 0 |
| 1 | 0 | 0 | 0 | 0 | 0 | 0 | 0 |
| 0 | 1 | 0 | 0 | 0 | 0 | 0 | 0 |
| 0 | 1 | 0 | 0 | 0 | 0 | 0 | 0 |
| 0 | 1 | 0 | 0 | 0 | 0 | 0 | 0 |
| 0 | 1 | 0 | 0 | 0 | 0 | 0 | 0 |
| 0 | 1 | 0 | 0 | 0 | 0 | 0 | 0 |
| 0 | 1 | 0 | 0 | 0 | 0 | 0 | 0 |
| 0 | 1 | 0 | 0 | 0 | 0 | 0 | 0 |
| 0 | 0 | 1 | 0 | 0 | 0 | 0 | 0 |
| 0 | 0 | 1 | 0 | 0 | 0 | 0 | 0 |
| 0 | 0 | 1 | 0 | 0 | 0 | 0 | 0 |
| 0 | 0 | 0 | 1 | 0 | 0 | 1 | 0 |
| 0 | 0 | 0 | 0 | 0 | 0 | 1 | 0 |
| 0 | 0 | 0 | 0 | 0 | 0 | 1 | 0 |

**Table S6.**
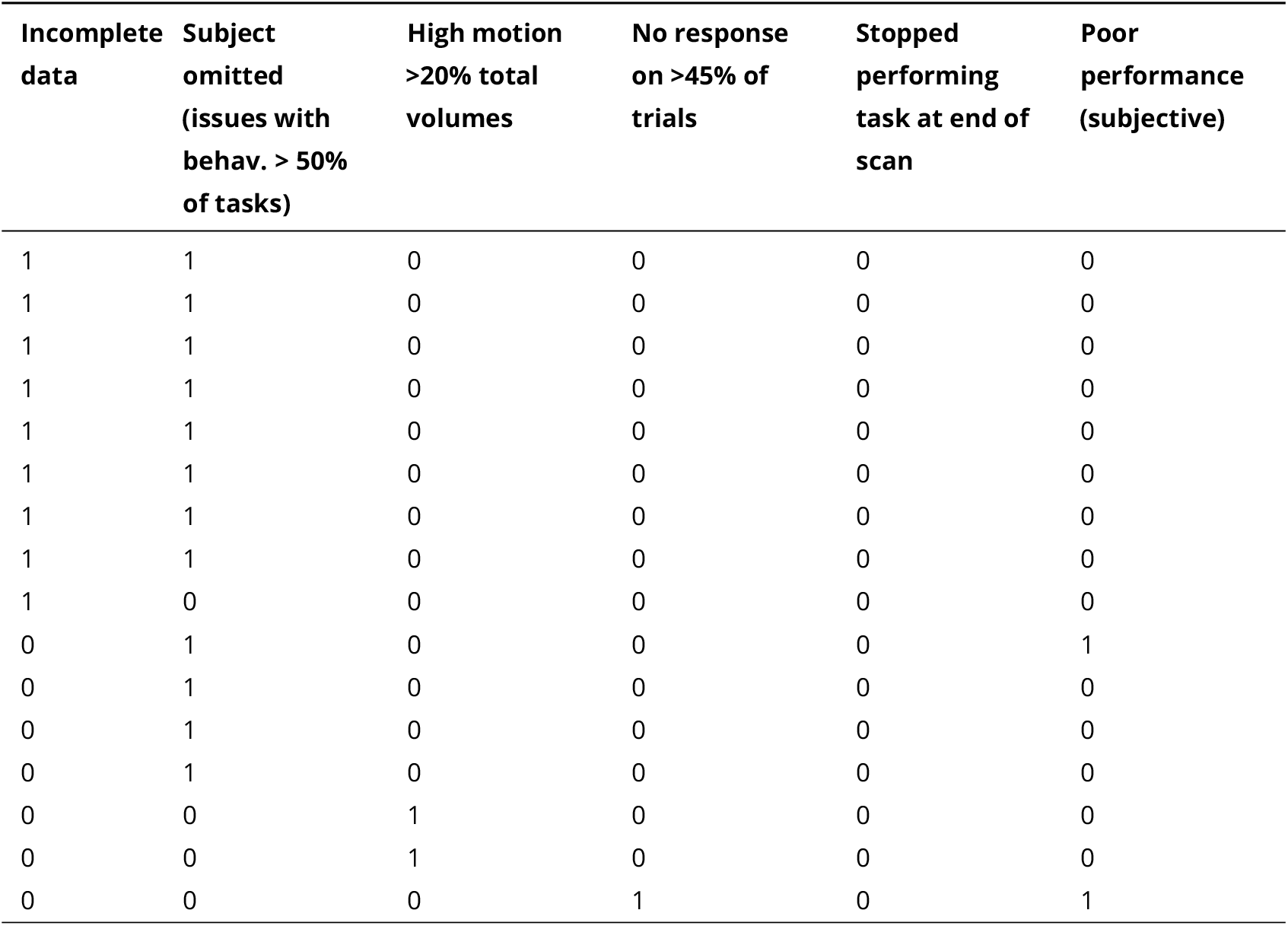
Exclusion information for Stroop task.

**Table S7.** Exclusion information for Cued Task Switching task.

| Incomplete data | Subject omitted (issues with behav. > 50% of tasks) | High motion >20% total volumes | No response on >45% of trials | Stopped performing task at end of scan | Poor performance (subjective) |
| --- | --- | --- | --- | --- | --- |
| 1 | 1 | 0 | 0 | 0 | 0 |
| 1 | 1 | 0 | 0 | 0 | 0 |
| 1 | 1 | 0 | 0 | 0 | 0 |
| 1 | 1 | 0 | 0 | 0 | 0 |
| 1 | 0 | 0 | 0 | 0 | 0 |
| 0 | 1 | 0 | 1 | 0 | 0 |
| 0 | 1 | 0 | 0 | 0 | 1 |
| 0 | 1 | 0 | 0 | 0 | 0 |
| 0 | 1 | 0 | 0 | 0 | 0 |
| 0 | 1 | 0 | 0 | 0 | 0 |
| 0 | 1 | 0 | 0 | 0 | 0 |
| 0 | 1 | 0 | 0 | 0 | 0 |
| 0 | 1 | 0 | 0 | 0 | 0 |
| 0 | 0 | 1 | 0 | 0 | 0 |
| 0 | 0 | 1 | 0 | 0 | 0 |
| 0 | 0 | 0 | 1 | 0 | 0 |

